# The Geometric Evolution of Aortic Dissections: Predicting Surgical Success using Fluctuations in Integrated Gaussian Curvature

**DOI:** 10.1101/2022.09.19.508582

**Authors:** Kameel Khabaz, Karen Yuan, Joseph Pugar, David Jiang, Seth Sankary, Sanjeev Dhara, Junsung Kim, Janet Kang, Nhung Nguyen, Kathleen Cao, Newell Washburn, Nicole Bohr, Cheong Jun Lee, Gordon Kindlmann, Ross Milner, Luka Pocivavsek

## Abstract

Clinical imaging modalities are a mainstay of modern disease management, but the full utilization of imaging-based data remains elusive. Aortic disease is defined by anatomic scalars quantifying aortic size, even though aortic disease progression initiates complex shape changes. We present an imaging-based geometric descriptor, inspired by fundamental ideas from topology and soft-matter physics that captures dynamic shape evolution. The aorta is reduced to a two-dimensional mathematical surface in space whose geometry is fully characterized by the local principal curvatures. Disease causes deviation from the smooth bent cylindrical shape of normal aortas, leading to a family of highly heterogeneous surfaces of varying shapes and sizes. To deconvolute changes in shape from size, the shape is characterized using integrated Gaussian curvature or total curvature. The fluctuation in total curvature (***δK***) across aortic surfaces captures heterogeneous morphologic evolution by characterizing local shape changes. We discover that aortic morphology evolves with a power-law defined behavior with rapidly increasing ***δK*** forming the hallmark of aortic disease. Divergent ***δK*** is seen for highly diseased aortas indicative of impending topologic catastrophe or aortic rupture. We also show that aortic size (surface area or enclosed aortic volume) scales as a generalized cylinder for all shapes. Classification accuracy for predicting aortic disease state (normal, diseased with successful surgery, and diseased with failed surgical outcomes) is **92.8 ±1.7%**. The analysis of ***δK*** can be applied on any three-dimensional geometric structure and thus may be extended to other clinical problems of characterizing disease through captured anatomic changes.

## 1 Introduction

Imaging modalities such as computed tomography (CT) provide sophisticated three-dimensional representations of the human body and are ubiquitously relied upon in clinical practice [1–4]. At its core, imaging captures anatomic changes that are linked to underlying pathologic processes. Anatomy is the spatial organization of tissues in space. As such, geometry is the natural mathematical framework to quantitate anatomy. Geometric approaches have widely been used to characterize and classify multiple diseases including pulmonary nodules [1], liver cirrhosis [2], and thyroid masses [3, 4]. In all of these examples, the physician’s human eye defines the hallmark of disease by the appearance of nodularity or spiculation. Even the process of aging or healing often leads to the appearance of undulations or wrinkles, which is a departure from the baseline smooth geometry of healthy skin [5]. These examples highlight the ubiquitous classification of anatomic objects based upon the surfaces that define their boundaries. These surfaces are easily appreciated when the anatomy is directly visualized with gross inspection (such as skin or with dissection in the operating room). They also appear as level-sets in cross-sectional imaging, most often x-ray based tomography, because of the intrinsic difference in x-ray absorption and scattering of various tissues. The problem of disease progression mathematically becomes the transformation of a smooth surface into a rough surface, with the appearance of multiple new length scales.

Quantifying what the human eye so easily discerns has proven challenging. Recent work has sought to translate successes from deep learning (DL) to utilize imaging for disease diagnosis and clinical planning [6, 7]. Challenges in the availability of high-quality data [8, 9], the lack of reproducibility and generalizability [9, 10], the disconnect between technical accuracy and clinical efficacy [11, 12], and interpretability of inherently “black-box” DL models have made the application of DL in medicine particularly fraught [16, 19]. Sophisticated approaches harnessing the machinery of Riemannian geometry have been successful in certain neuroanatomy problems [17]; however, they are limited to tissues with well defined internal landmarks allowing for global coordinate systems, e.g. Talairach space or MNI coordinate system in brain imaging [18], which do not exist for cardiovascular tissues.

We take a fundamentally mathematical approach to developing imaging-based geometric descriptors of disease evolution in clinically meaningful systems. We focus on the challenging problem of classification in aortic dissections. The aorta is the largest blood vessel in the human body. It functions as a conduit carrying blood from the left ventricle of the heart to provide blood flow to the head, arms, abdominal organs, and legs [20, 21]. Mechanically, the aorta is akin to a pressurized distensible cylindrical shell; aortic pathologies are mechanical in nature, with aortic rupture leading to near certain death. Aortic dissections are partial tears in the aortic wall. Mechanically, dissections are cracks which do not penetrate through the entire aortic thickness; rather, they propagate between the different concentric layers making up the aortic wall generating aortic blisters (medically termed the false lumen). Dr. Michael DeBakey, the pioneer of aortic dissection surgery, famously described diseased thoracic aortas as ‘torturous’, ‘globular’, ‘U-shaped’, ‘dilated’, and ‘idiosyncratically varying’ [22]. These qualitative descriptors underpin the complex changes in shape that accompany aortic disease and its morphologic evolution. The current standard of care for type B aortic dissection (TBAD) is anchored on the concept of preventing “dangerous” shape changes, such as aneurysmal degeneration, which contribute to worse outcomes [21, 23]. Thus, the concept of shape and pathology are well-accepted clinically; however, accurate quantification of these changes is limited. Thoracic aortic stabilization by placement of a fabric-covered stent into the diseased thoracic aorta, clinically termed Thoracic Endovascular Aortic Repair or TEVAR, is the preferred modern surgical approach to TBAD [21, 23].

Since the 1950’s and DeBakey’s initial work, the ubiquitous anatomic classifier clinically used to trigger aortic surgery has been maximum aortic diameter (2*R*_*m*_, where *R*_*m*_ is maximum radius) [21, 23], yet clinicians have long appreciated that size by itself is an inadequate descriptor of aortic anatomy [23–26] and have resorted to qualitative descriptors of aortic morphologic evolution. The urgent need for a richer description of aortic anatomy has been highlighted by several clinical trials using TEVAR, which showed that using size alone to trigger intervention did not improve mortality [27–30]. The cardio-vascular surgical community realizes that better identification of patients who would benefit from TEVAR is needed to shift the post-intervention mortality curve in the positive direction [29–32]. Identifying anatomic parameters that impact the ability of an aorta to remodel [33] is instrumental in guiding therapy.

Geometry and mechanics have driven the use of maximum diameter as a singular scalar descriptor of aortic stability and, therefore, the dominant trigger of intervention. Geometrically, since aortas are generalized bent cylinders, the cylinder’s radius naturally appears as a principal length scale describing the geometry. Since diseased aortas expand non-uniformly in the radial direction as a function of axial distance (or distance along the aortic centerline), the gradient of aortic radius along the centerline is maximized near the largest diameter. Clinically, largely for the ease of application, the gradient has been replaced by simply measuring the maximum diameter and assuming a uniform ‘normal’ aortic radius. This assumption is fraught with problems because normal aortic diameters, while being uniform for any given individual, show a broad distribution in the general population [34]. The problem is further complicated by the fact that uniform aortic growth (∼0.5 mm/year) occurs with aging [35]. Lacking appropriate normalization, the maximum diameter is only a descriptor of Egeometry in comparison to population means.

Standard biomechanics literature asserts that stability of dilated aortas is dominated by the Law of Laplace, where the stress (*σ*) is a simple linear function of internal pressure (*P*), aortic thickness (*h*), and maximum diameter (2*R*_*m*_): *σ* = *P* ·*R*_*m*_/*h*. By Laplace, larger aortas will have linearly proportional larger stresses. Since aortic mechanical stability is a fracture problem, once the wall stress supersedes a critical stress, the aorta will crack. Given the distribution of *R*_*m*_ values in the population, ⟨*R*_*m*_⟩ + δ*R*_*m*_ ∼1.25 ±0.5 cm, and the fact that *R*_*m*_ is not constant with time, δ_*t*_ ⟨*R*_*m*_⟩ = 0.5 mm/year, makes calculation of a broadly applicable critical stress nearly impossible. Furthermore, the mathematical derivation of Laplace’s law assumes homogeneous isotropic shells of uniform thickness, which is not true even for non-diseased aortas [13]. More advanced mechanical failure analysis of aortic aneurysms and dissections utilizes finite element approaches (FEA); however, these ultimately lack the simple relationships between geometry (derived from clinical imaging data) and stress contained in Laplace’s law. As such, their wide clinical application has been limited [14, 15]. Nearly all clinical data gathered about aortas is imaging-based containing geometric information. The herculean efforts to translate intricate mechanical stability models using FEA to clinical practice largely failed due to lack of interpretability and understanding of these models by practicing physicians. This failure should be heeded when developing sophisticated AI-based deep-learning models which take as inputs raw CT scan data; the challenge of interpretability of such ‘black-box’ models in medical applications is well known [16, 19].

We derive and validate a mathematical descriptor of aortic shape; a single variable that scales appropriately with DeBakey’s famous qualitative descriptors of aortic shapes [22, 24]. As D’Arcy Thompson wrote a century ago: “In a very large part of morphology, our essential task lies in the comparison of related forms rather than in the precise definition of each” [36]. The problem of morphogenesis is deeply linked to the stability of self-reproducing structures; Rene Thom postulated that any morphological process is divided into islands of determinism, dominated by morphogenic fields whose precise mathematical form is defined by topologic homeomorphisms, separated by regions of instability and indeterminacy [37]. Topology is now a fundamental tool in the classification of rapidly changing shapes and structures in developmental biology, where anatomic descriptions are translated into topological language allowing the quantification of purely geometric transformations in growing tissues [38–40]. The generalization of a tissue or body into sets of smooth, closed, orientable surfaces (i.e. membrane systems) makes them topological objects [37, 40]. Local topological surgeries, cutting or gluing, lead to global topological changes in biological forms through regions of topological catastrophes [37, 39, 40]. We apply such methods to the study and generalization of aortic morphogenesis throughout the life-cycle of an aorta (see figure 1).

**Fig. 1.**
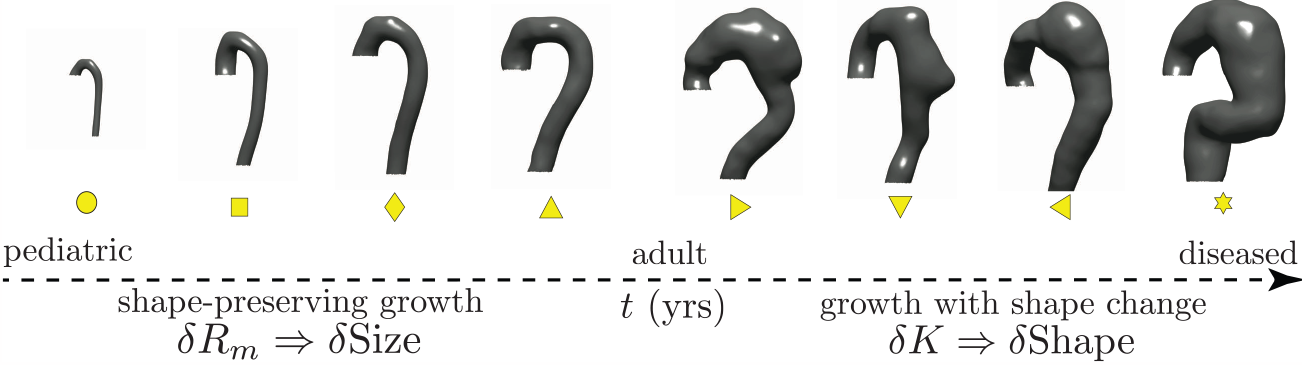
Eight representative aortas along the normal-to-diseased axis (left to right): a 1-year-old child, healthy adults, and type B aortic dissection (TBAD) patients at varying degrees of aneurysmal degeneration. Two clinical regimes exist: shape preserving growth and growth with shape changes.

We work with aortic surfaces derived from cross-sectional imaging such as CT scans from both normal patients and patients with aortic dissections at various stages of disease. Figure 1 outlines the problem of defining a continuous geometric parameter space that can smoothly evolve from a normal pediatric aorta to the most diseased aortic dissection. Visual inspection of the objects in figure 1 easily leads to a qualitative description of their anatomy: the aortas grow left to right with an initial rapid change followed by a more subtle increase in size (scale), a trend that reverses when surface shape is considered. The rapidly growing aortas are self-similar concerning their shape (bent cylinders with the toroidal aortic arch followed by the cylindrical descending thoracic aorta); a shape symmetry-breaking transition occurs as the aorta further moves towards the right. We follow this transition by projecting the CT-derived anatomy of 302 aortas, distributed from normal to diseased states, into a novel geometric space spanned by orthonormal axes for shape and size defined using the shape operator and total curvature of the aortic surface. The local geometry of the aortic surface is defined by its principal curvatures, k_1_ and k_2_. Specific measures of aortic shape in the literature have focused on functions of mean (*κ*_*m*_ = (*k*_1_ + *k*_2_)/2) or Gaussian (*κ*_*g*_ = *k*_1_*k*_2_) curvature [41, 42], cross-sectional deviations [43], and calculations on centerline reductions of the aortic surface [25, 44, 45]. A mathematically robust and reproducible definition of aortic shape does not exist despite these efforts because of spatial over-reduction and convolution of shape and size information. Centerline calculations lose information when the aortic surface deforms heterogeneously [46], and *κ*_*g*_, the dominant shape function, proves itself an ill-defined shape measure because it convolutes shape and size changes [46]. We solve this problem by using the Gauss-Bonnet Theorem and the total curvature, *K* = ∬_*A*_ *κ*_g_*dA*, as the primary measure of shape. We prove that all aortic shapes are homeomorphic to **T**^2^; as such, all aortas no matter how deformed remain generalized bent cylinders. Using simulated shapes with evolving surface roughness, we show that *κ*_*g*_ and K indeed hold the same shape information content, provided overall changes in size are small.

Topology preservation is by far not the norm in biologically growing surfaces that are non-conservative and can locally add mass or remove mass without the constraint of elastic deformations or linkage to some global manifold [footnote 1]. The existence of a homeomorphism implies that as aortas deform, be it through normal growth or pathology, every increase in K in some region must be balanced by a proportional decrease somewhere else on the aortic surface. Therefore, the distribution of *K* across the aortic surface holds information about shape. This allows us to classify shape by studying the statistical properties of these distributions. Since ⟨*K*⟩ is constant, the variance of *K* captures the balance of positive and negative curvature regions across the surface: *δK* = ⟨*K*^2^⟩ − ⟨*K*⟩ ^2^. *δK* captures the heterogeneous morphologic evolution by characterizing local shape changes. We hypothesize and demonstrate that this novel shape measure correlates with aortic disease evolution and can be used to characterize treatment response for aortic dissections when appropriately coupled with a size metric.

## 2 Methods

### 2.1 Clinical Data Cohort

We analyzed a cohort of 302 computed tomography angiography (CTA) scans from 2009 and 2020 (171 of non-pathologic aortas (Appendix A.1 figure 13) and 131 scans of diseased aortas (Appendix A.1 figure 14)). In the diseased cohort, two pathologies were included: TBAD (124 scans) and thoracic aortic aneurysms (7 scans). Three patient subgroups are derived from the main cohort to analyze geometric predictors of outcomes following TEVAR: a control cohort, a failed TEVAR cohort, and a successful TEVAR cohort. The control cohort consists of 171 scans of non-pathologic aortas (Appendix A.1). Failed repair is defined as reintervention or type IA endoleak in the follow-up period (using the most recent patient data available). [Footnote 2]

A balanced cohort of 18 patients with desired TEVAR outcomes and 18 patients with failed outcomes is analyzed. Patients all met the inclusion criteria of having both a pre-operative and post-operative CTA. A total of 45 scans (18 pre-operative, 27 post-operative) are included for the desired outcomes group and 51 scans (18 preoperative, 33 post-operative) are included for the failed outcomes group. Outcomes are defined by the presence or lack of need for secondary surgical intervention more than 30 days from index TEVAR. Given the retrospective nature of this data cohort, all decisions to reintervene were made by the primary surgeon. Most common reasons for reintervention post-TEVAR were continued false lumen (FL) expansion and type 1A (proximal seal zone) leak. Demographic, disease, and imaging information for this group is summarized in Appendix A.1. All data collection and analysis is performed in accordance with the guidelines established by the Declaration of Helsinki and under institutional review board approval (IRB20-0653 and IRB21-0299).

### 2.2 Meshing Algorithm and Geometric Parameterization

Three-dimensional aortic models are created from CTA image data using a custom workflow in Simpleware ScanIP (S-2021.06-SP1, Synopsys, Mountain View, CA). Segmentation of the aortic geometry from the scans is processed using a five-step algorithm which includes 1. segmentation, 2. noise reduction, 3. smoothing, 4. isolation of the outer surface, and 5. surface meshing. A representative schematic of the process is shown in Figure 2 and more information on the process can be found in Appendix A.2. A triangular mesh for the outer surface is generated for each smoothed segmentation in ScanIP for analysis in Matlab (2021b, Mathworks, Natick, MA). A total of 15 meshed surfaces are generated for each segmentation (sampling 5 mesh densities and 3 smoothing variations), allowing for control of process-derived variance in surface curvature calculations.

**Fig. 2.**
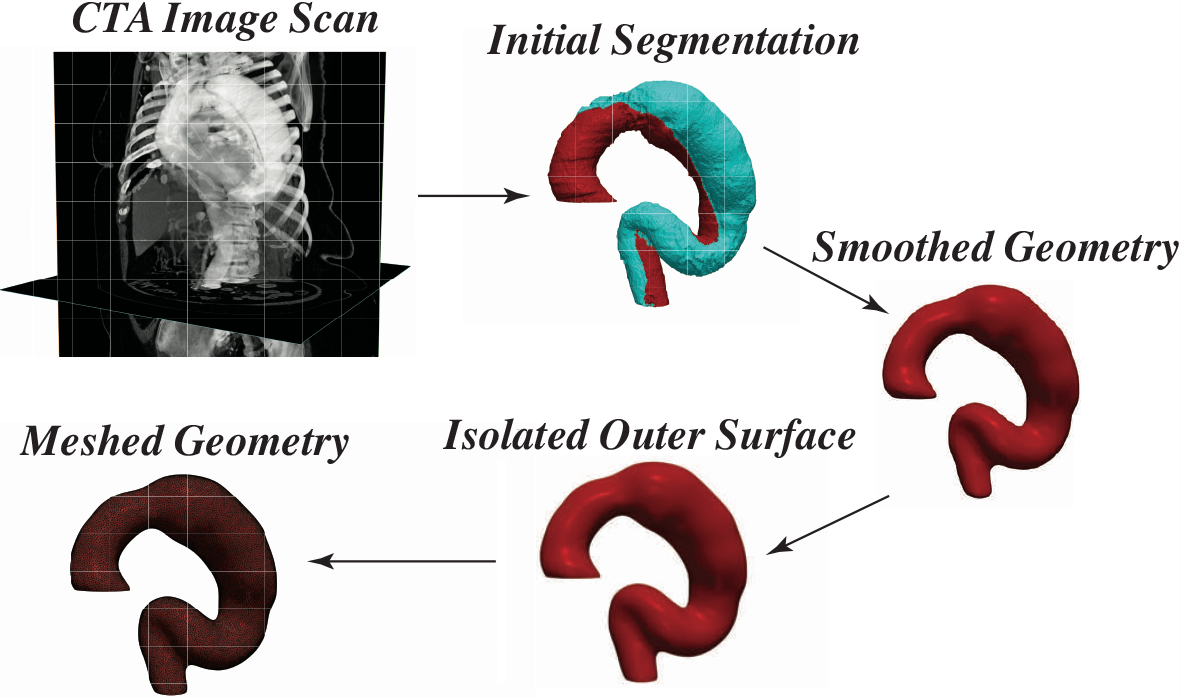
Aortic Segmentation and Post-Processing Workflow. The aorta is segmented from a chest CTA imaging scan. The segmentation is then smoothed and the outer surface is isolated. A mesh is then placed on the geometry for surface analysis.

Once a meshed geometry is created for each aorta, the Rusinkiewicz algorithm is used to calculate the per-vertex shape operator 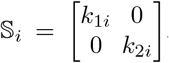, where *k*_1*i*_ and *k*_2*i*_ are the per-vertex principal curvatures. Briefly, the algorithm calculates the per-vertex shape operator as a weighted average of the shape operators of immediately adjacent faces. The per-face tensors are computed using a finite-difference approximation defined in terms of the directional derivative of the surface normal [50] (see Appendix A.3 for more details). Therefore, each vertex in the mesh is accompanied by a first and second principal curvature that approximate the local shape at the intersection of neighboring faces. Appendix A.4 outlines the artifact removal process that was implemented.

### 2.3 Size and Shape Characterization

#### 2.3.1 Size

Figure 3 shows our computational workflow for calculating aortic shape and size features. Aortic size is parameterized using the centerline length *ℒ* and aortic radius *ℓ*. While *ℒ* is singular for each aorta, *ℓ* is a distribution especially in diseased aortas with heterogeneous shapes, and therefore a family of radii. Multiple measured values can be used to represent this distribution to characterize aortic size: mean aortic radius (⟨ *R* ⟩), median aortic radius 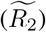, and maximum aortic diameter 2*R*_*m*_. *ℓ* can also be calculated directly from surface curvatures. The mean Frobenious norm of the per-vertex shape operator 𝕊_*i*_, 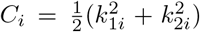, is the per-vertex Casorati curvature [47–49]. Averaging over the entire aorta gives the mean Casorati curvature,

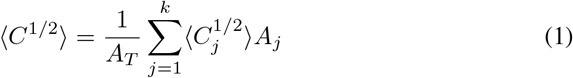

where 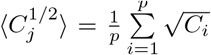 is the per-partition mean, *k* is the number of partitions, *A*_*j*_ is the area of a partition, and p the number of vertices within each partition. The need for dividing the surface into *k* smaller partitions is not necessary to obtain accurate mean Casorati curvatures; however as explained below, it becomes necessary when performing shape calculations. We keep it here from an implementation consistency standpoint because *C*_*i*_ is calculated from principal curvatures. The inverse mean Casorati curvature ⟨*C*^1*/*2^⟩ ^*-*1^ is a measure of aortic size along the direction of greatest curvature, which is the aortic radius. Lastly, higher order functions of size such as total aortic area, *A*_*T*_, and aortic volume, *V*, are calculated directly from the surface segmentations. Total aortic area is also calculated from the sum of per ele-ment areas: 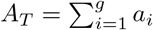, where g is the total number of triangular mesh elements on a given aortic surface. It is important to note that only ⟨*C*^1*/*2^⟩ ^*-*1^ allows for a meaure of aortic size using only information contained in 𝕊_*i*_; all other size measures necessitate additional information.

**Fig. 3.**
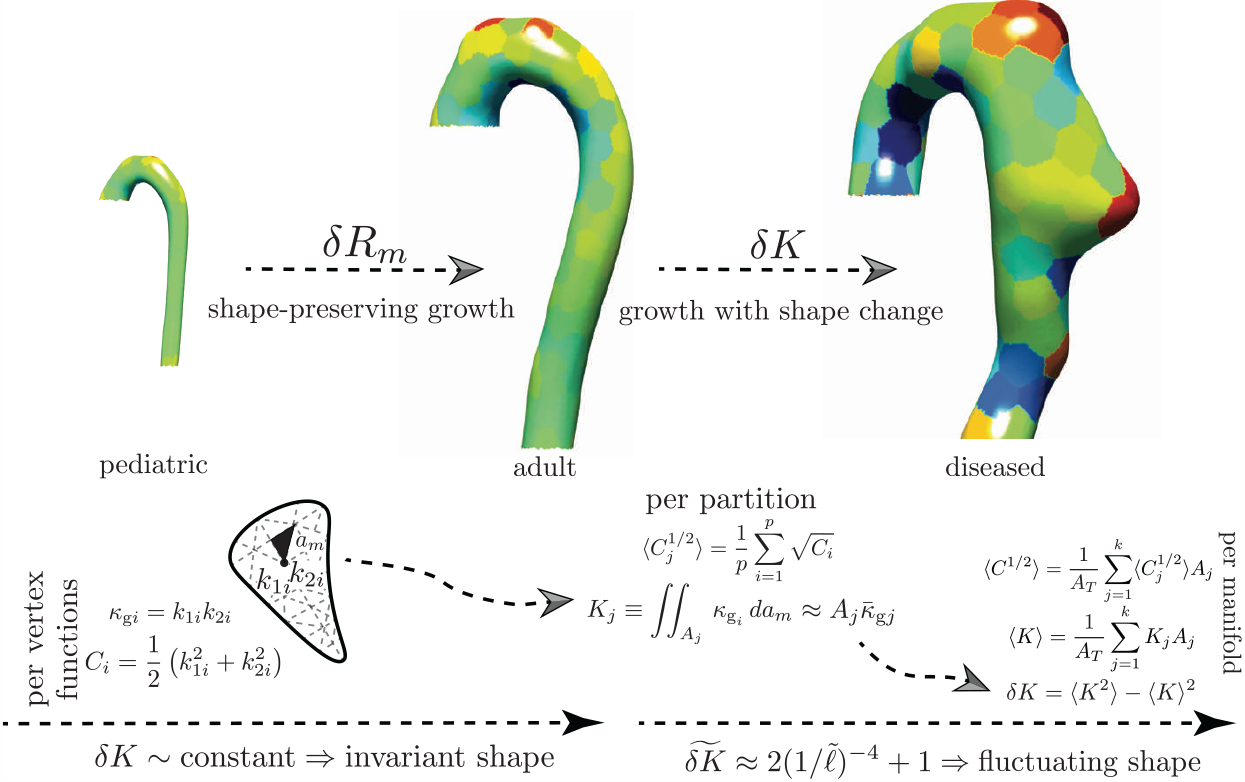
By mapping the aortic surface to the unit sphere (Gauss map) [46], we have an independent measure of shape. The per-vertex shape operator 𝕊_i_ is calculated using the Rusinkiewicz algorithm [50]. To minimize noise, the aorta is divided into multiple partitions with area *A*_*j*_. The local integrated Gaussian curvature *K*_*j*_ is calculated as the product of each partition area and mean Gaussian curvature, 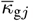 is equivalent to the signed partition area 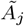 mapped out by the normals projected onto the unit sphere. We define aortic shape by studying the statistics of the distributions of *K*_*j*_. ⟨*K*⟩ and *δK* are the first and second distribution moments that define aortic shape geometry, respectively.

#### 2.3.2 Shape

The size-independent parameterization of shape is obtained through an effective mapping of normals on the external aortic surface *𝒮* to the unit sphere **S**^2^. Operationally, we do not directly perform the Gauss mapping. Instead, we calculate the total curvature over local regions (partitions) of the aortic surface with area A_*j*_, which by the local Gauss-Bonnet theorem is the holonomy angle of the circumnavigated area, a topologic quantity [47]. In our analysis, given geometrically highly heterogenous shapes, a meshed surface is employed and the per vertex Gaussian curvature is extrinsically calculated from the shape operator: *κ*_*gi*_ = |𝕊_*i*_| = k_1*i*_k_2*i*_. The area elements are intrinsic to the surface and 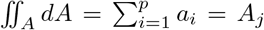; however, since a generic triangular mesh is used, no information about the surface metric exists. Knowledge of the metric would be needed to carry out the integration of per-vertex Gaussian curvature explicitly. Our computational approach therefore relies on splitting the integration into a product 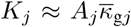,where 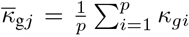. To bring the mean Gaussian curvature within a partition outside the integral, *κ*_*gi*_ must vary weakly within *A*_*j*_. Clearly, this variance is minimized when *A*_*j*_ = *a*_*i*_. As shown in Appendix A.6 (see Figure 17 panel IV), calculating *K* at this scale holds little information about overall aortic shape. To understand the necessity of sub-dividing the aortic surfaces into partitions where *a*_*j*_ < *A*_*j*_ < *A*_*T*_, the scale space nature of the problem needs to be explored and appreciated. Since the per-vertex Gaussian curvature is calculated on a mesh with an average element spacing proportional to the CT scan resolution (∼ 𝒪 (0.5) mm); this value assumes the mesh-imposed inner scale. However, as is appreciated by examining the evolution in Figure 1, the change in aortic shape, the increasing ‘bumpiness’, is occurring on a much larger length scale. Visual inspection of the shapes shows this scale to be on the order of the aortic radius or *ℓ*. The mesh-imposed inner scale is ∼ *ℓ* /1000. As discussed in Appendix A.2, the mesh-resolution was set by the inherent resolution of the CT data and by the need of a finely enough spaced computational mesh to calculate discrete derivatives of surface normals. However, from an aortic shape classification standpoint, using the mesh-imposed inner scale runs the risk of imposing a large amount of “spurious resolution” or “false detail”, commonly seen with scale space problems when an inappropriately small inner scale is selected [47]. We impose *ℓ* as the inner-scale by sub-dividing the aortic surface into k area partitions via a Voronoi decomposition using the k-means algorithm, in which *k* = *A*_*T*_ / *ℓ* ^2^ (see figure 4 which shows that k is independent of how *ℓ* is measured and relatively constant with respect to *ℓ*). For subsequent analysis, 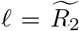 (indicating the median *R*_2_, where *R*_2_ = 1/*k*_2*i*_ is calculated per-vertex from 𝕊_*i*_). We use the k-means++ algorithm to find initial centroid seeds, and the k-means algorithm is performed with a maximum of 10000 iterations. Appendix A.6 shows how *k* can vary by a factor of 10 and does not impact the results discussed in the following sections. Lastly, Appendix A.5 outlines our use of the Jensen-Shannon Divergence to check that at this inner-geometric scale Gaussian curvatures still show acceptably small variance, allowing us to bring the mean Gaussian curvature within a partition out of the area integral and replace the area integral with the sum of mesh element areas.

**Fig. 4.**
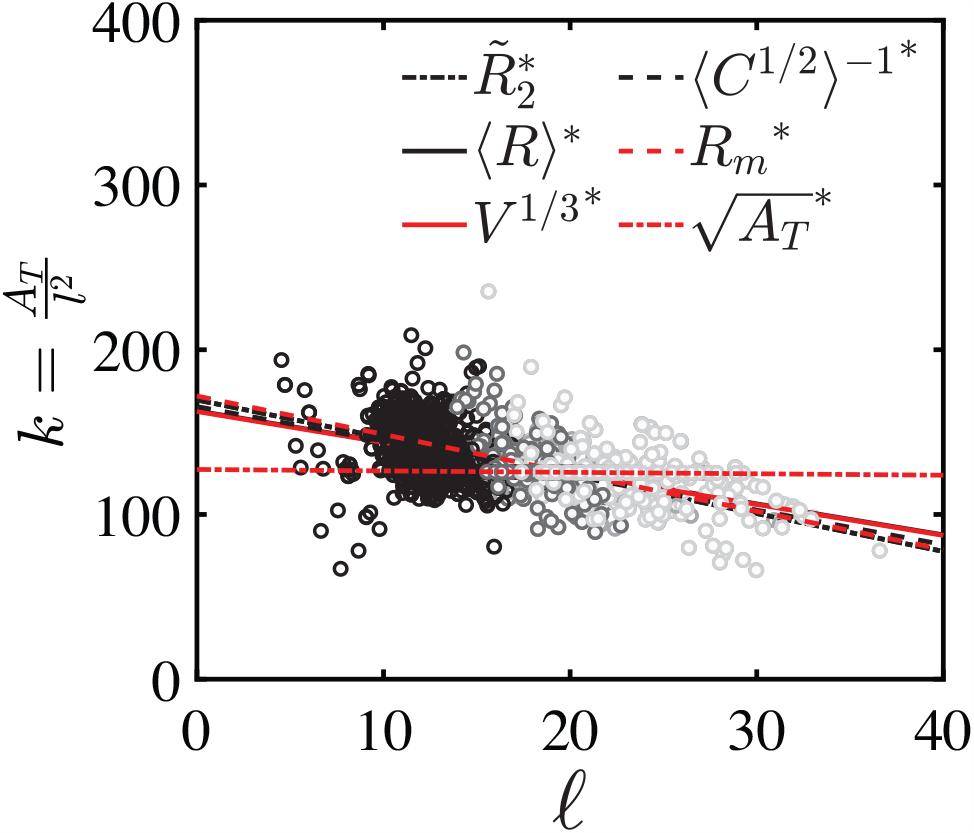
Data for 302 aortas, including non-pathologic (black circles), pathologic with failed TEVAR (light gray circles), and pathologic with successful TEVAR (dark gray circles) aortas are plotted. The linear scaling can be used to define *A*_*j*_ ∼ *ℓ*^2^, which sets the number of partitions *k* used in the Gauss map calculations. The various linear fits are taken for different definitions of size outlined in section 2.3.1. In this case, the fits are normalized by the pre-factors from figure 5 collapsing the data.

Having partitioned the aortic surface into k partitions of size ∼ *ℓ* ^2^, the per partition total curvature

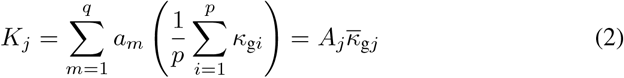

is calculated. To characterize the entire aorta, the sum of per partition total curvatures (a topologic invariant)

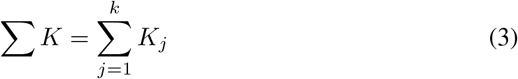

and the fluctuations in total curvature (a normalized measure of shape) across the entire manifold surface are calculated:

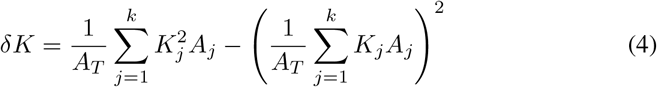

The mean Gaussian curvature

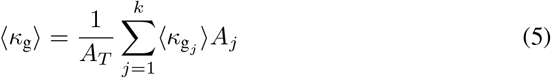

and the fluctuation in Gaussian curvature of the manifold

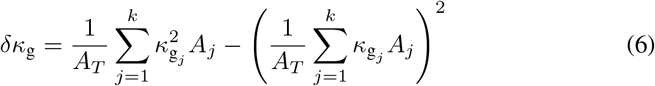

are also calculated.

#### 2.3.3 Statistics

Each aorta contains k partitions. All calculations are performed for each of the 15 meshed models per aorta. Of note, the k-means partitioning is independently performed for each mesh. An average value for ⟨*C*^1*/*2^⟩, Σ*K, δK*, ⟨*κ*_*g*_⟩, and δ*κ*_*g*_ is computed for the 15 meshes, such that each scan becomes represented by a single value that aggregates the variation from the smoothing and meshing algorithms. To quantify variability from the partitioning procedure itself, the entire process (partitioning, partition-level curvature calculations, manifold-level curvature calculation, and averaging between meshes) is repeated 10 times to obtain 10 replicates of each value per aorta. The mean of these replicates is reported and a sensitivity analysis is performed in Appendix A.6. The error bars in figure 7 represent ±1 standard deviation to quantify the variability in the results from the partitioning replicates.

### 2.4 Aortic Feature Space Classification using Machine Learning

The classification accuracy of different shape and size metrics in determining aortic disease states (normal aortas, successful TEVAR, and failed TEVAR) is calculated using the means of the distributions, the mean of the means of the distributions, and a logistic regression classification. The distribution mean-based methods model is reflective of current clinical practice, in which clinicians are most cognizant of the characteristics of a “typical” patient in each group (the mean) and are less aware of the variation within each group (the standard deviation). The first model defines each threshold t as the mean value of the parameter for the two neighboring distributions. After these boundaries are defined, each scan is assigned a “predicted” classification according to its geometric characteristic, and the accuracy of the predictions is computed. The second model defines each threshold as the mean value of the means of the two neighboring distributions.

A multinomial logistic regression with 1000 random permutations of train-test splits with a 50% training and 50% testing distribution is used for the third model. Logistic regression is a classification method that models *p*(*X*) = *P r*(*Y* = 1| *X*), the probability that some response variable *Y* takes on a specific value of 1 (a binary classification) based on the input data *X*, using the logistic function: 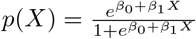

The parameters *β* _0_ and *β* _1_ are estimated from training data using the maximum likelihood method. The accuracy of this model in predicting binary outcome *Y* is determined by subdividing the dataset into a training set and a testing set, fitting *β*_0_ and *β*_1_ using the training set, and computing the error rate between model predictions and data for the testing set.

As the classification problem is between three classes — non-pathologic aortas, diseased aortas with desired outcomes following TEVAR, and diseased aortas with failed outcomes following TEVAR — multinomial logistic regression is used. Multinomial logistic regression is an extension of logistic regression for the setting of H > 2 classes with coefficient estimates defined as follows:

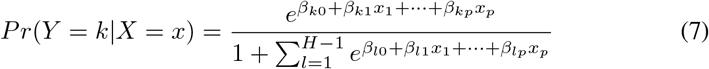

The variability of the logistic decision boundary includes lasso regularization applied to ⟨C^1*/*2^⟩ to examine the impact of increasing the model’s dependence on shape over size. The objective function is the penalized negative binomial log-likelihood:

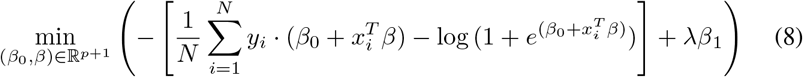

Two decision boundaries are created using two binomial logistic regression classifiers. The hyperparameter λ is varied to understand the impact of decreasing the influence of size on clinical decision-making to maximize the impact of using a shape-based classifier [52, 53].

### 2.5 Finite Element Simulations

The utility of K over *k*_*g*_ is investigated in a more controlled setting by simulating local growth in ideal geometries and examining the associated change in *k*_*g*_ versus K. Two geometries are tested: a sphere that experiences a small change in global size followed by surface deformation due to local growth and an idealized aorta that experiences a larger change in global size followed by surface deformation due to local growth. Growth is modeled using a morphoelastic model in ABAQUS (2018, Dassault Systèmes, Waltham, MA) that decomposes the deformation gradient F into an elastic contribution F_*e*_ and a growth contribution F_*g*_ : F = F_*e*_F_*g*_ [54–56]. Note that the Neo-Hookean (NH) strain energy function is only computed based on the elastic contribution. All parameters are determined from F_*e*_ instead of the total deformation F in the case growth is triggered. When there is no growth, F_*g*_ = I and F_*e*_ = F, which represents a purely elastic deformation. With this decomposition and the NH strain energy, the stress can be derived and updated during the loading process according to the following relations:

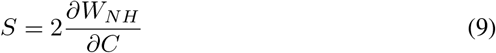

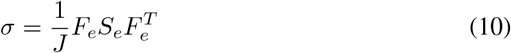

Where S_*e*_ and *σ* are the second Piola-Kirchhoff stress and Cauchy stress, respectively. The above model is implemented with ABAQUS Explicit solver and using the VUMAT subroutine [57, 58]. Assuming isotropic growth with constant growth rate in the longitudinal-circumferential plane, the growth contribution F_*g*_ becomes:

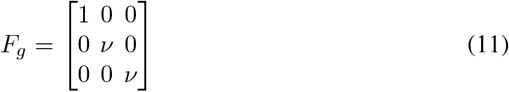

where *v* is the growth factor and the growth rate is 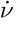. See Appendix A.9 for further details.

## 3 Results

There are three major and two minor results that come from our analysis. The first major result (3.1) is that aortic size scales as a generalized bent cylinder and depends only on a single length scale, *ℓ*. The second major result (3.2) is that projecting aortas into a two-dimensional space defined by (*δK, ℓ* ^*-*1^) separates aortic geometries into shape-preserving self-similar growth and growth with shape changes; moreover, all aortic geometries are shown to be topologically homeomorphic to **T**^2^. The third major result (3.4) shows that the (*δK, ℓ* ^*-*1^)-feature space outperforms all other size and shape measures when applied to the predictive classification of TEVAR success. The first minor result (3.3) shows that the data projected into the normalized 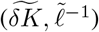 space are best fit to a simple power law 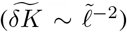. The second minor result (3.5) shows that *δK* and *δK*_*g*_ hold similar information, however, *δK* provides a stronger signal of shape changes when shape and size changes occur simultaneously.

### 3.1 Universal Scaling of Aortic Size

The analysis includes a variety of size measures previously described in the literature, including the traditional metric of maximum diameter (2*R*_*m*_) and higher-dimensional values of area (*A*_*T*_) or volume (V) (figure 5). When each length scalar is plotted versus maximum diameter, the data collapses onto lines, with linear size measures having a slope of 1 and higher-dimensional measures (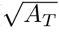 and *V* ^1*/*3^) having slopes that correspond to modeling aortic size as a generalized bent cylinder. In generalized bent cylinders, area and volume scale as *A*_*T*_ ∼ 2π *ℓ* × *ℒ* and *V* ∼ π *ℓ* ^2^× *ℒ*. Hypothesizing a linear relationship between axial length *ℒ* and cross-sectional circular radius *ℓ, ℒ* ∼ c*ℓ*, the reparameterized equations *A*_*T*_ ∼ 2π c *ℓ* ^2^ and V ∼ π c *ℓ* ^3^ are obtained.

**Fig. 5.**
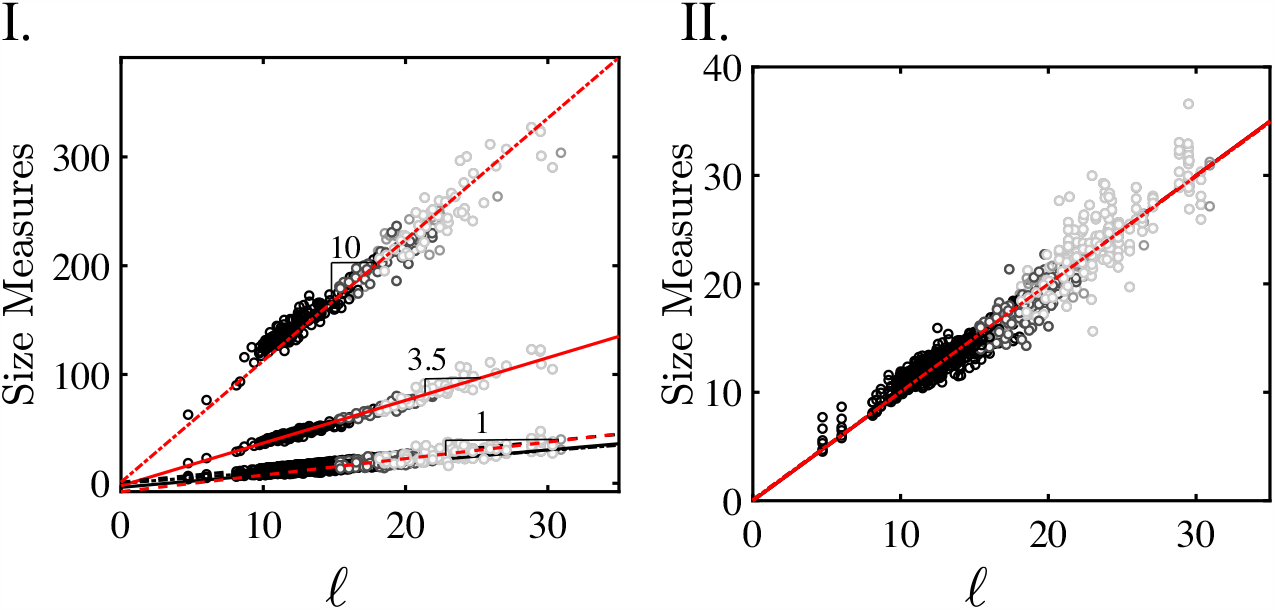
Data for 302 aortas, including non-pathologic (black circles), pathologic with failed TEVAR (light gray circles), and pathologic with successful TEVAR (dark gray circles) aortas are plotted. I. shows that parameterizations of aortic size (mm) including maximum aortic diameter (2*R*_*m*_, red dashed line), mean radius (⟨*R*⟩black solid line), median radius (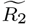, black dotted line), and mean inverse linearized aortic Casorati curvature (⟨*C*^1*/*2^ ⟩^*-*1^, black dashed line) are equivalent. Dimensionally scaled, aortic area 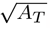 red dotted line) and volume (*V* ^1*/*3^, red solid line) are also linear when plotted against *ℓ* = 2*R*_*m*_. All size measures can be collapsed onto a single master curve (II.), proving that all aortas scale as generalized bent cylinders parameterizable by a single length scale *ℓ*.

Figure 5 shows that 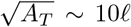 and *V* ^1*/*3^ *∼* 3.5 *ℓ*, which implies that *A*_*T*_ *∼* 100 *ℓ* ^2^ and *V ∼* 42.88 *ℓ* ^3^. Solving for c, we obtain c *∼* 15.9 for area and c *∼* 13.6 for volume. The per-factor c can alternatively be calculated using *ℒ ∼* c *ℓ* directly. *ℒ* is best approximated using the length of the aortic centerline measured from the segmentations. Thus, 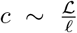 can be calculated on each aortic geometry (Figure 6), with *c* = 16.6 *±*2.4 when *ℓ* ∼ ⟨*C*^1*/*2^ ⟩is used. The values of *c* ∼15.0, *c* ∼13.6 and *c* ∼16.6 indicate similar constants that are produced using independent methods and that quantify a linear relationship between aortic axial length and cross-sectional size. This demonstrates that aortic geometry follows a universal linear scaling between different size measures and globally retains an invariant toroidal-cylindrical geometry. This indicates that additional information is unlikely to exist with higher-dimensional size measurements that have been the focus of recent literature and provides geometric reasoning behind the well-known utility of maximum diameter in aortic management.

**Fig. 6.**
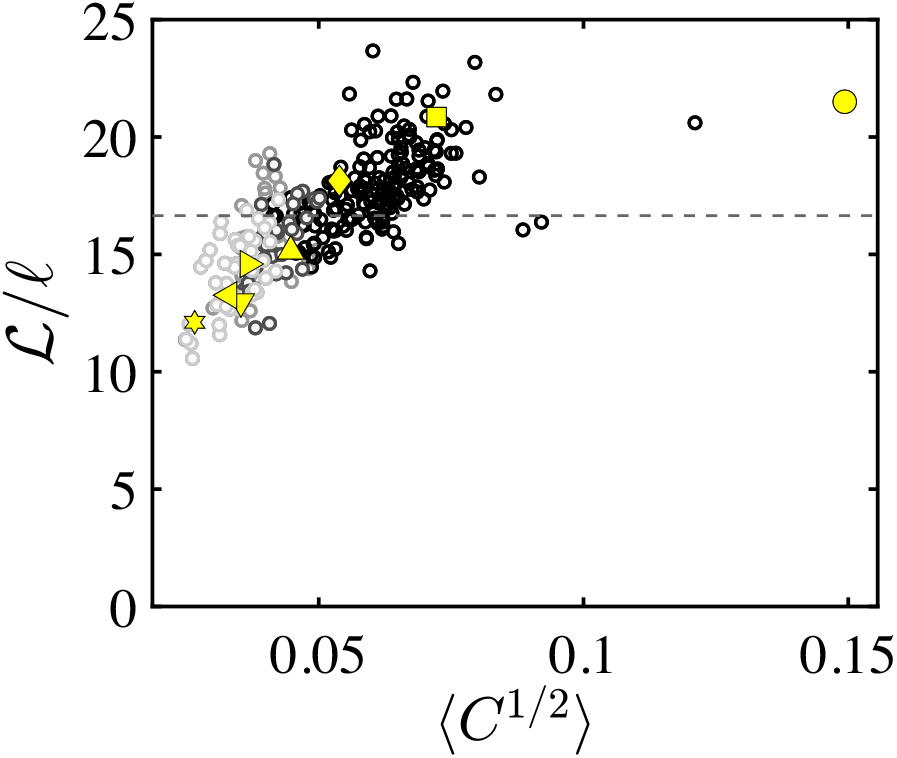
Ratio of centerline length *ℒ* to radial size *ℓ*. For the relationship *ℒ ∼ c ℓ* indicating a linear scaling between axial length and cross-sectional circular radius, we obtain *c* = 16.6 *±* 2.4.

**Fig. 7.**
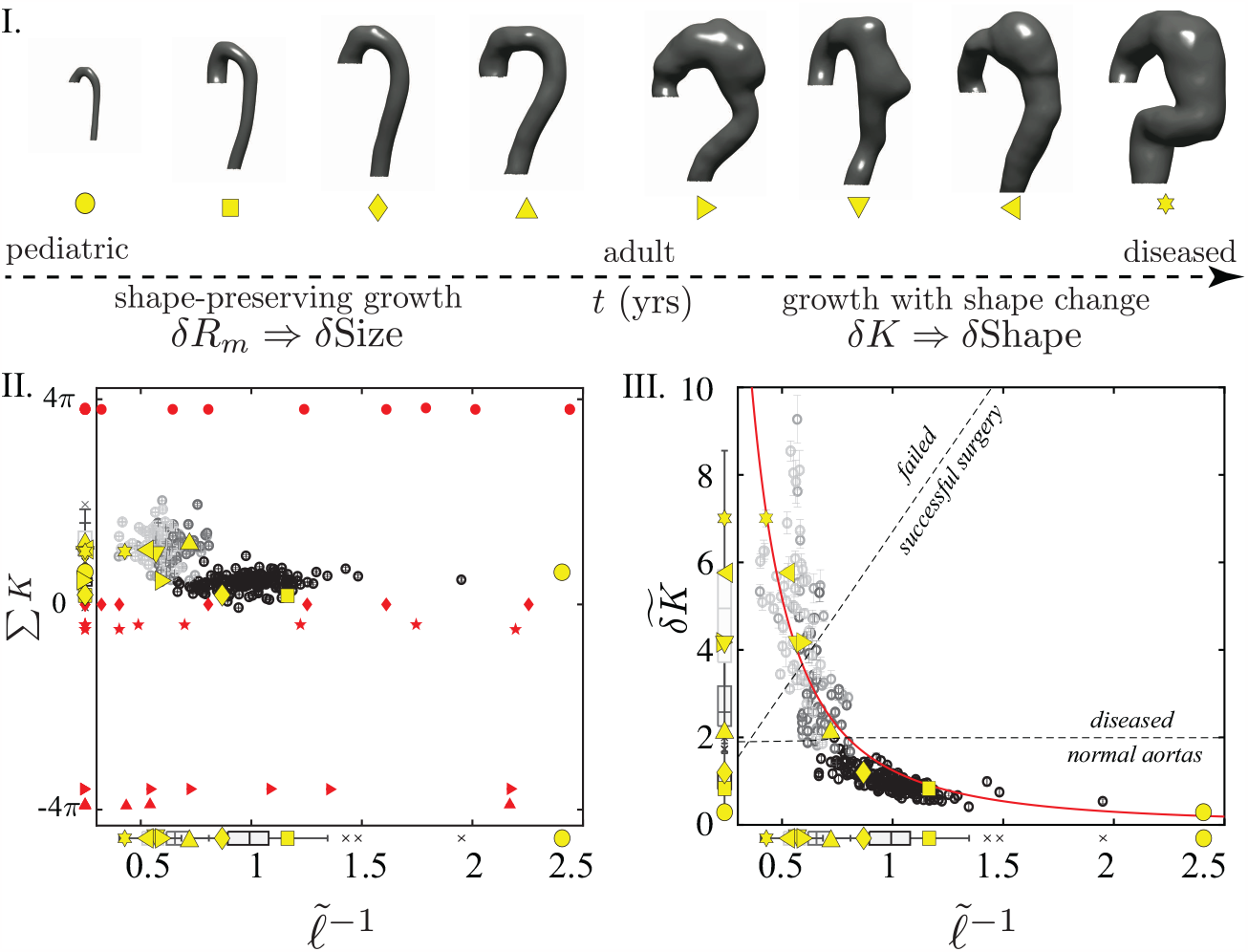
I. The eight canonical representative aortas along the normal-to-diseased axis (left to right): a 1-year-old child, healthy adults, and type B aortic dissection (TBAD) patients at varying degrees of aneurysmal degeneration. Two clinical regimes exist: shape preserving growth and growth with shape changes. II. shows the topologic equivalence of all aortic shapes to tori (red stars) and cylinders (red diamonds); the yellow symbols correspond to specific aortic shapes along the normal-to-diseased axis (I.). Red circles correspond to perfect spheres of varying size; pseudospheres and catenoids are depicted as red rightward-pointing triangles and upward-pointing triangles, respectively. III. shows the optimal two-dimensional aortic geometric feature space with independent axes for size and shape. The solid red curve 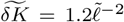 is a best fit to the data. The power-like behavior is further supported by the probability distribution of *bK* (figure 8). The aortas separate into shape invariant (normal) and shape fluctuating (diseased) populations. Furthermore, this feature space defines decision boundaries which correctly classify diseased patients based upon success of TEVAR.

### 3.2 Role of Shape Deviation

Figure 7 plots the dat from 302 aortic segmentations into two spaces: 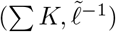 and 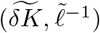. Size is represented in both by 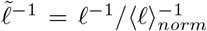, where ⟨ *ℓ* ⟩_*norm*_ is the mean of the normal aortas. As was shown in 3.1, the different measures of ℓ differ only by a pre-factor, which is constant throughout the data. Therefore, the same normalized size axis is obtained irrespective of how *ℓ* is measured. Σ*K* is the total integrated Gaussian curvature of each aorta and mathematically defines its topology. Figure 7II plots the aortic data points (black and grey circles) along with the representative aortas in yellow. Red points are calculated on analytically generated surfaces (see Appendix A.7) analyzed through the same analysis pipeline as the aortic data: spheres (red circles), tori (red stars), cylinders (red diamonds), pseudospheres (red right-ward pointing triangle), and catenoids (red upward pointing triangle) of various sizes. These ideal shapes represent the three canonical geometries: spherical, parabolic, and hyperbolic. Their total curvature values of (spherical), Σ*K* = 0 (parabolic or Euclidian), and Σ*K* = −4π (hyperbolic) are in agreement with analytical results from differential geometry [59]. This agreement for the ideal surfaces is an important validation of the computational methodology used in this paper. The aortic data clearly align along the Σ*K* = 0 line and cluster with the tori and cylinders. The normal aortas (solid black dots) show a very tight distribution with very little variance along Σ*K*. The diseased aortas show more spread, which is likely a consequence of the complex shapes encountered in the group. Nevertheless, what is striking in this data is that throughout the morphologic life span of an aorta (during normal growth and diseased degeneration), any given aorta is topologically equivalent to a generalized bent cylinder or torus homeomorphic to **T**^2^. Consequently, by the Gauss-Bonnet Theorem, aortic topologic invariance implies that as aortas deform, be it through normal growth or pathology, every increase in *K* somewhere must be balanced by a proportional decrease elsewhere on the aortic surface [46, 51, 59]. Therefore, the distribution of K across the aortic surface should hold information about shape. Since ΣK is constant, the variance *δK* = ⟨*K*^2^⟩ ⟨*K* ^2^⟩ captures the balance of positive and negative curvature regions across the surface and is a measure of shape deformation. Normalized δ*K* is defined as 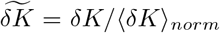, where ⟨ *δK* ⟩_*norm*_ is the mean *δK* of the normal aortas. Examination of the data in figure 7 III shows that 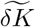 quantitatively captures the above-mentioned qualitative descriptions of aortic shape. Our three aortic populations naturally separate into three regions in 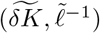-space. In the asymptotic limit 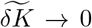, self-similar aortic growth occurs; all normal aortas exist along this limit. In the other limit, 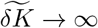, fluctuations in shape become seemingly independent of size change. The most distorted and convoluted appearing aortas, which also correspond to patients with failed endovascular surgeries, cluster in this limit. The two limits are joined through a transition region that appears as an elbow in the two-dimension shape-size feature space. Interestingly, the dissection patients who had successful surgery predominantly cluster in this region. The utilization of this novel 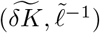-space for aortic classification is further developed in section 3.4.

### 3.3 Power Law Fit in the 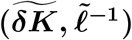 Feature Space

In the 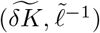 geometric feature space, the data can be fit to a power law 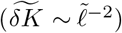. Here we follow an established statistical framework of discerning power-law behavior in empirical data to determine whether the data is truly consistent with a power-law [61]. Figure 8 shows separate probability distributions for δ*K* and *R*_*m*_ and then plots these on doubly logarithmic axes. Logarithmic distributions, *p*(*x*) *x*^*-α*^, are linear in log-log space: ln *p*(*x*) = α ln *x* + *c*, where *c* is a constant. The log-log transformed *δK* distribution suitably fits a straight line with *R*^2^ ≈ 0.74, while the *R*_*m*_ data does not fit a line (*R*^2^ ≈ 0.002). This indicates that a power law may be an appropriate fit for the shape metric δ *K* and is not an appropriate fit for the size parameter *R*_*m*_. The size data are well approximated by a Gaussian distribution, a fact demonstrated in the literature on aortic size distributions [34].

**Fig. 8.**
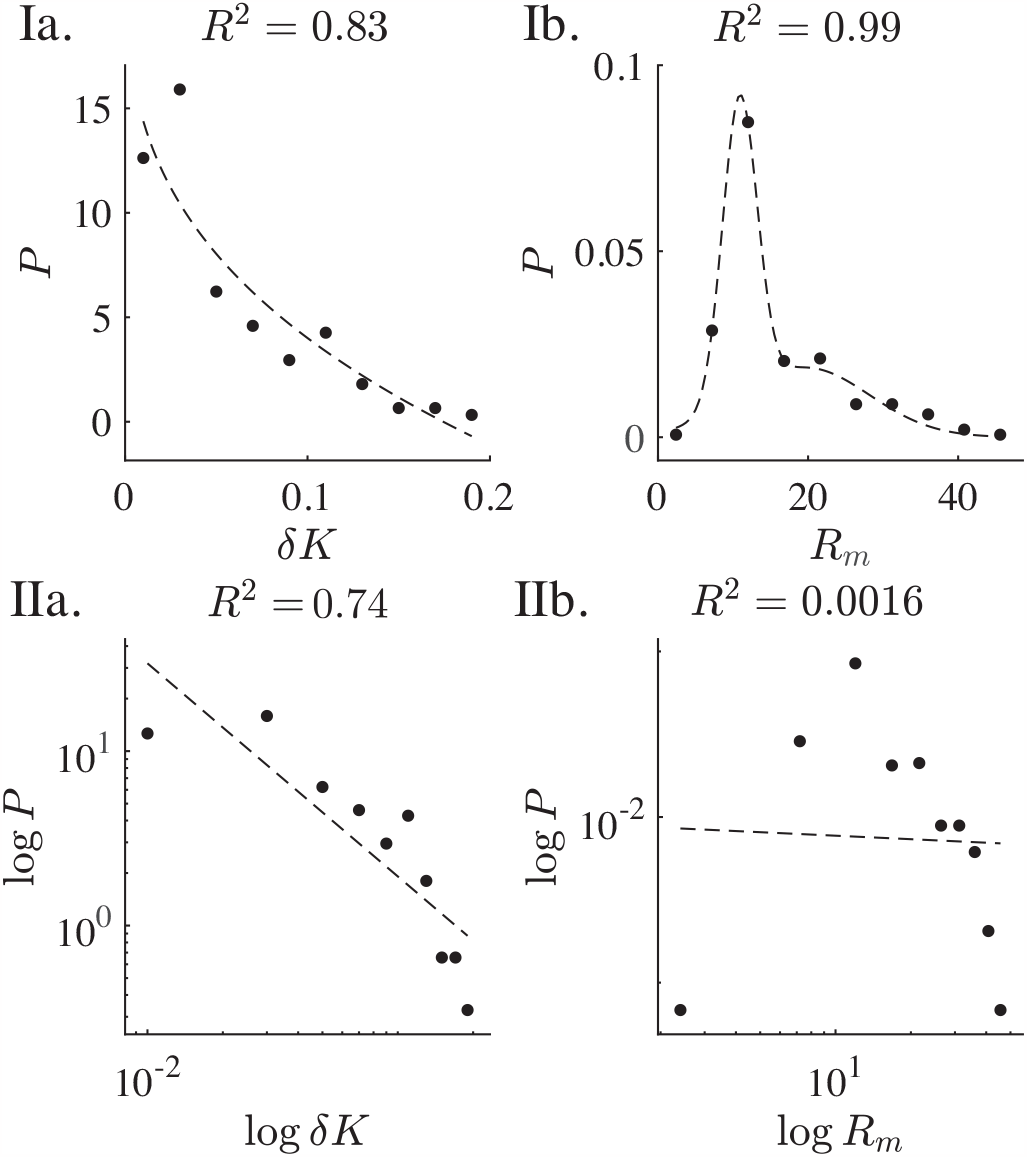
Probability distributions of *bK* and *R*_*m*_ are plotted. Ia. The *bK* distribution is fitted to a power law in the form *P* = *ax*^*b*^ + *c* IIa. A linear fit log *P* = *b* log *bK* + *c* achieves a high *R*^2^. Ib. The *R*_*m*_ distribution is well-fit to a two-term Gaussian in the form 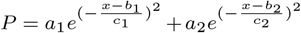 IIb. When a linear fit is applied to the log-transformed data, log *P* = *b* log *R*_*m*_ + *c*, a low *R*^2^ value results.

### 3.4 Classification with Predictive Modeling

The strong correlation between *δK* divergence and the evolution of aortic pathology begs application to treatment planning. Figure 9 compares the effectiveness of aortic size, the clinical standard, with the 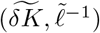 shape-size feature space in determining aortic disease states (normal aortas, successful TEVAR, and failed TEVAR). Significant variability in aortic size within the broader patient population is captured by this single institutional dataset. Furthermore, there is overlap between the normal and diseased populations and as such, classification accuracy dramatically changes due to slight differences in model parameterizations (from 73% in figure 9 Ia, 83.9% in figure 9 Ib, and 87.0 *±* 2.3% in figure 9 Ic). The dual parameterization of both aortic size and shape — two orthogonal metrics — expands the geometric characterization into a two-dimensional space (figure 9 II). Classification accuracy increases to 90.3% in figure 9 IIa and 92.8 *±* 1.7% in figure 9 IIb.

**Fig. 9.**
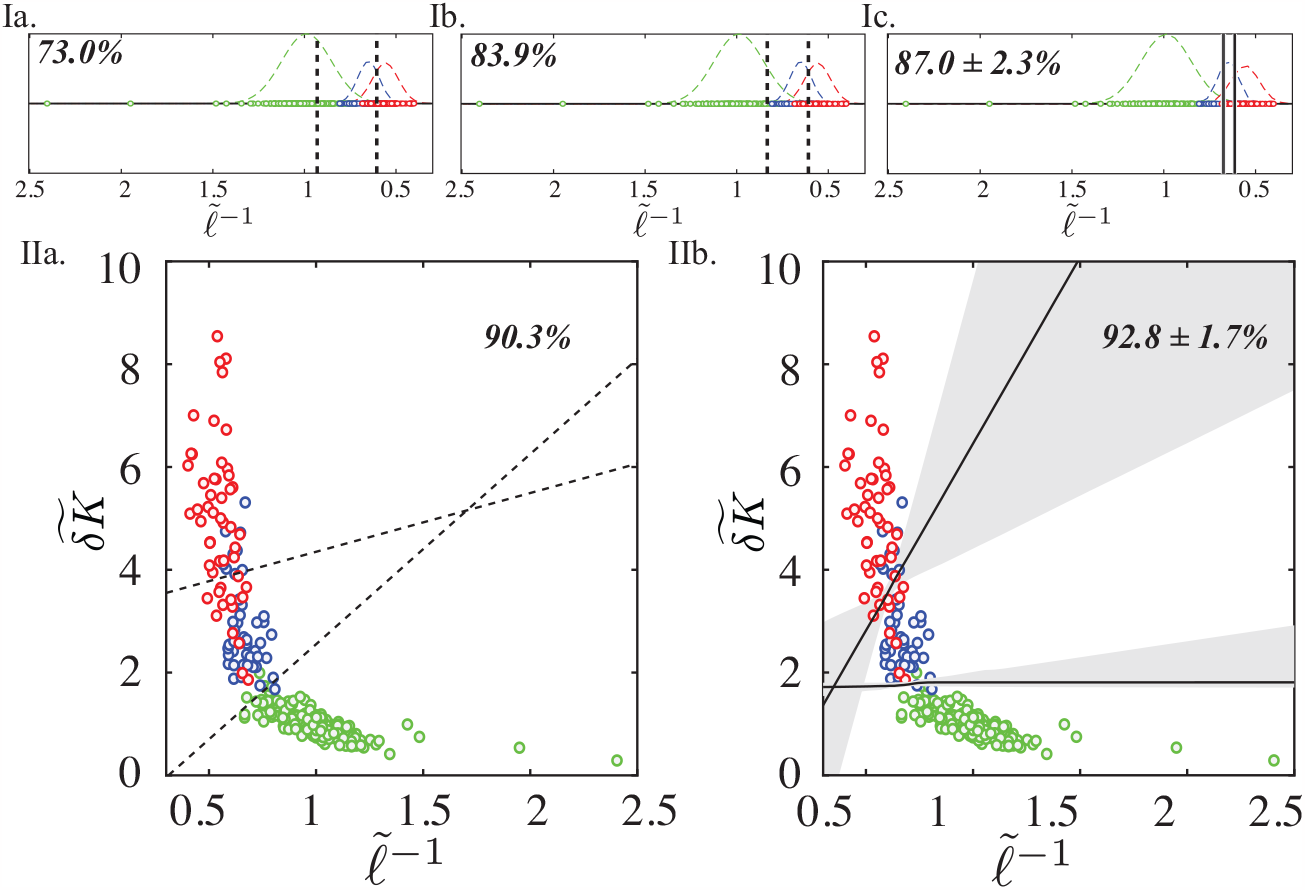
The 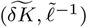 geometric feature space improves upon current sized-based methods. I. The clinical paradigm relies on size metrics alone to classify aortic disease states (green for normal aortas, blue for successful TEVAR, and red for failed TEVAR). However, broad within-group size distributions indicate the considerable variability in aortic sizes within the general population. Clinicians routinely utilize statistical means of these distributions as thresholds for classifying disease states, but linear decision boundaries are highly sensitive to small changes in model setup. Ia. A 73.0% accuracy for classifying the 3 states is obtained when each threshold is defined as the mean ⟨ *C*^1*/*2^ ⟩ of the two neighboring distributions. Ib. An 83.9% accuracy is achieved when the threshold is defined as the midpoint of the means of individual class distributions. Ic. A 87.0% mean accuracy is obtained when a logistic regression classifier is used. Thus, small changes in how a threshold is defined dramatically alter the perceived utility of size. II. The 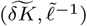 shape and size-based geometric feature space allows for the utilization of two independent parameters to characterize aortic disease state. IIa. A 90.3% classification accuracy is obtained when defining thresholds according to the mean *bK* and ⟨ *C*^1*/*2^ ⟩ of each patient group. IIb. A 92.8% mean accuracy with a standard deviation of only 1.7% is obtained using a logistic regression classifier with varying regularization parameter. The shaded region indicates the interquartile range of decision boundaries and demonstrates the robustness of the two-parameter space. Unlike the single parameter space, the presence of two physically interpretable and orthogonal asymptotic limits ensures more effective classification.

As shown in Fig. 10-Ia, there is no significant difference amongst size predictors such as aortic volume, surface area, median and maximum diameters shown in figure 5, as well as Gaussian-curvature based size measures like the L2-norm of the Gaussian curvature (GLN) and the area-averaged Gaussian curvature (GAA) [82– 84], see Appendix A.8. There is a major difference with shape measures: clinical shape measures, which are based on the aortic centerline and include tortuosity index [25, 44, 45, 88], question mark angle [89, 90], cross sectional eccentricity [43, 91], and mean centerline curvature [92], significantly underperform *δK* in aortic disease state classification (Fig. 10-Ib). Similarly, *δK* outperforms other shape measures described in the biomedical engineering literature including the sphericity index (*Χ*) [87], flatness index (*γ*) [87], Gaussian curvature (*κ*_*g*_) [41, 42], area-averaged mean curvature (MAA), and L2-norm of the mean curvature (MLN) [82–84].

**Fig. 10.**
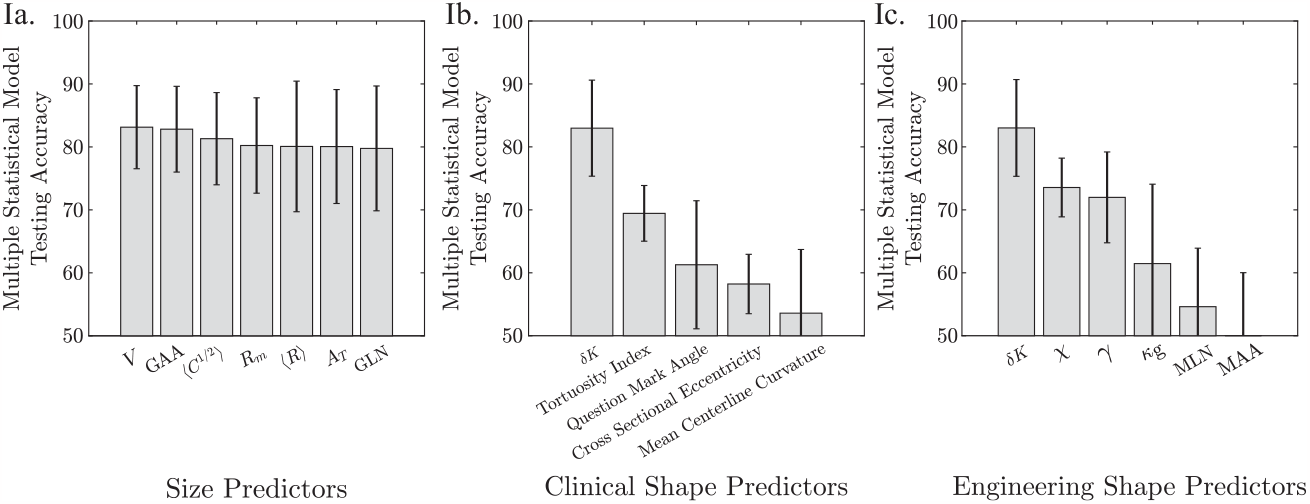
Comparison of size and shape metrics in classifying aortic disease state from medical imaging. Ia. Measures of aortic size achieve similar classification accuracies and thus are functionally equivalent (corroborating figure 5). The GLN and GAA are other size metrics, as shown in figure 18. *bK* significantly outperforms measures of aortic shape from the clinical literature in classifying aortic disease state (normal non-diseased aortas, diseased aortas with desired outcomes following TEVAR, and aortas with failed outcomes following TEVAR). Ic. *bK* outperforms general shape metrics from the broad engineering literature. Error bars indicate *±*1 standard deviation of the classification accuracies for the different classification methods.

### 3.5 Shape Evolution Modeling

In this paper, K is shown to be a topologic invariant across aortic anatomies and *δ K* a strong function of aortic shape changes. The integrated total curvature Σ *K* is a topologic invariant and remains approximately constant across all aortic anatomies (Figure 7 II). *δ K* is approximately constant across the range of normal aortas, matching empirical knowledge that non-pathologic aortas experience shape-preserving growth. *δ K* diverges with rapidly degenerating aortas, especially those where TEVAR ultimately failed. The role of *δ K* in this divergence is elucidated with FE simulations on spheres (Figure 11) and an idealized aorta (Figure 12). More information on the simulation parameterizations can be found in Appendix A.9.

**Fig. 11.**
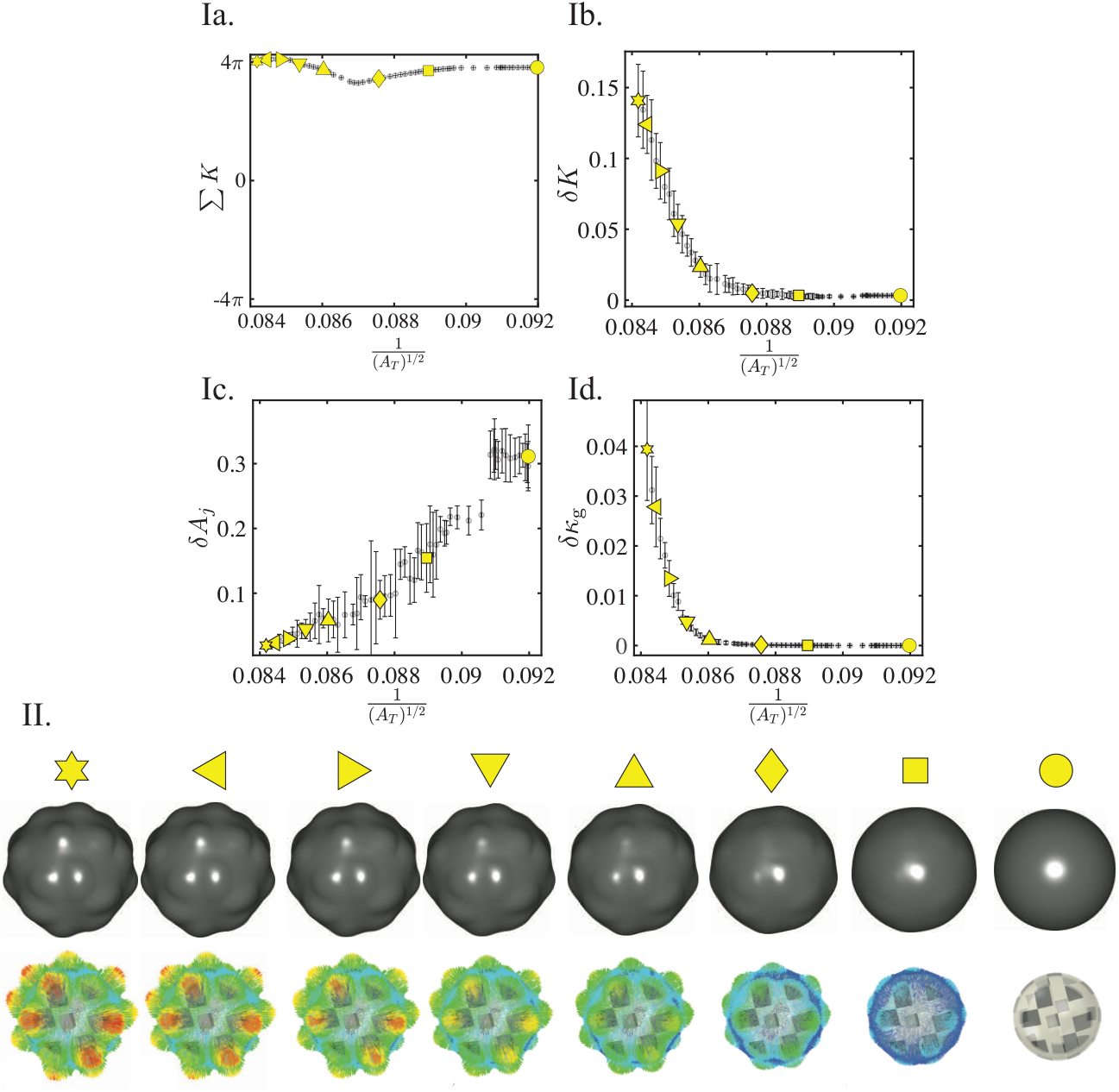
Simulation of a sphere with pressurization followed by growth. Ia. Σ*K* is a topologic invariant and thus remains relatively constant throughout the simulation. Ib. *δK* captures the increasing degeneration of the spherical surface due to growth. *Ic. δA*_*j*_ fails to capture this degeneration. Id. *δκ*_*g*_ seems to capture the degeneration of the spherical surface as the value diverges for increasing size. However, the narrow scale of the x-axis indicates that there is little increase in overall size for this simulation. II. Surface geometries for selected frames in the simulation, with the undeformed geometry on the right side and the final geometry on the left side. The vectors indicate the direction of surface deformation.

**Fig. 12.**
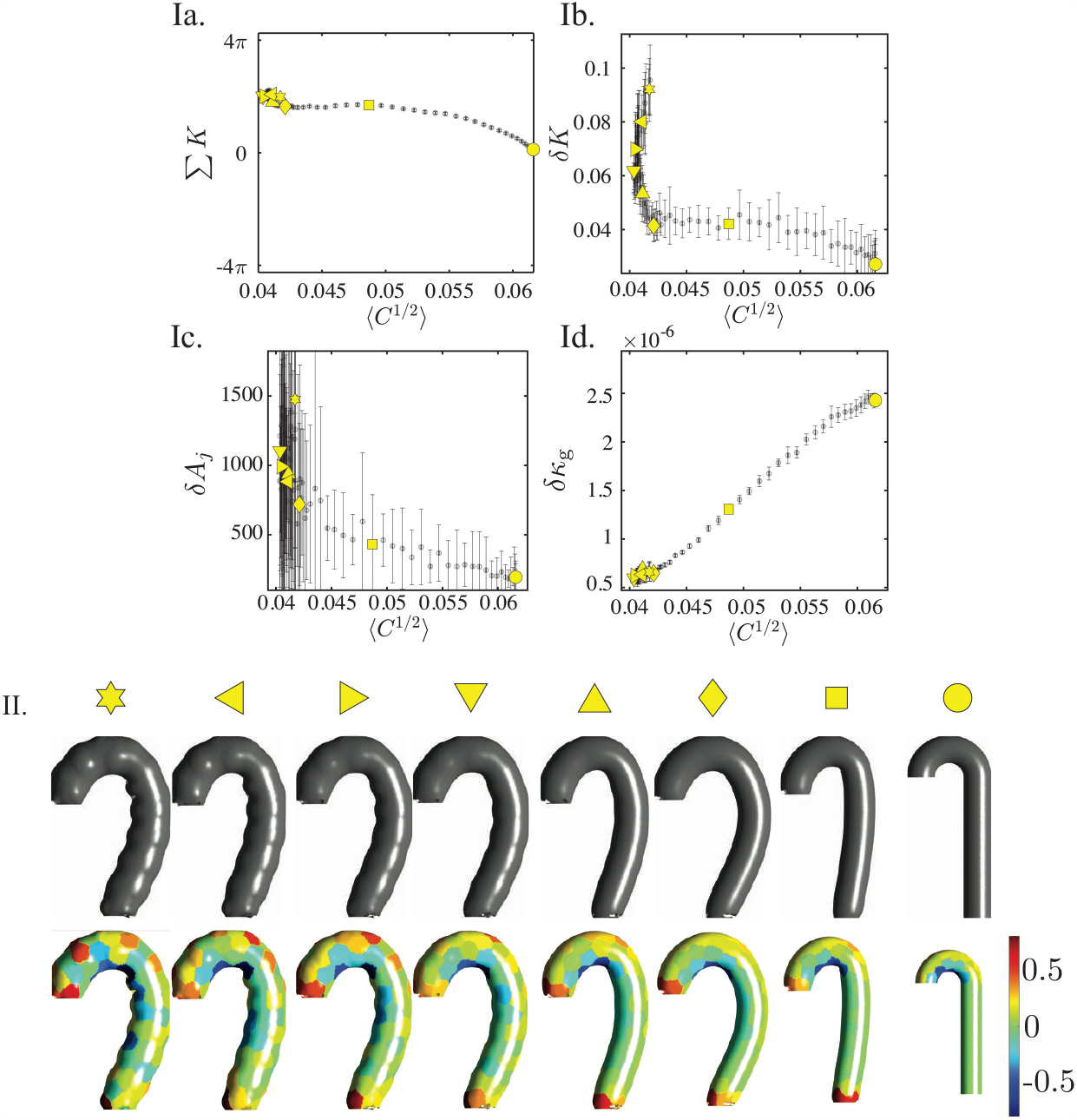
Simulation of an ideal aorta with pressurization followed by growth. Ia. Σ*K* is a topologic invariant and thus remains relatively constant throughout the simulation. Ib. *δK* captures the increasing surface degeneration due to growth. Ic. *δA*_*j*_ does not capture this degeneration. Id. When size significantly changes, *δκ*_*g*_ no longer captures the geometric deformation. II. Surface geometries for selected frames in the simulation, with the undeformed geometry on the right side and the final geometry on the left side. The heatmap coloring indicates *K*_*j*_ the total curvature at the per-partition level.

**Fig. 13.**
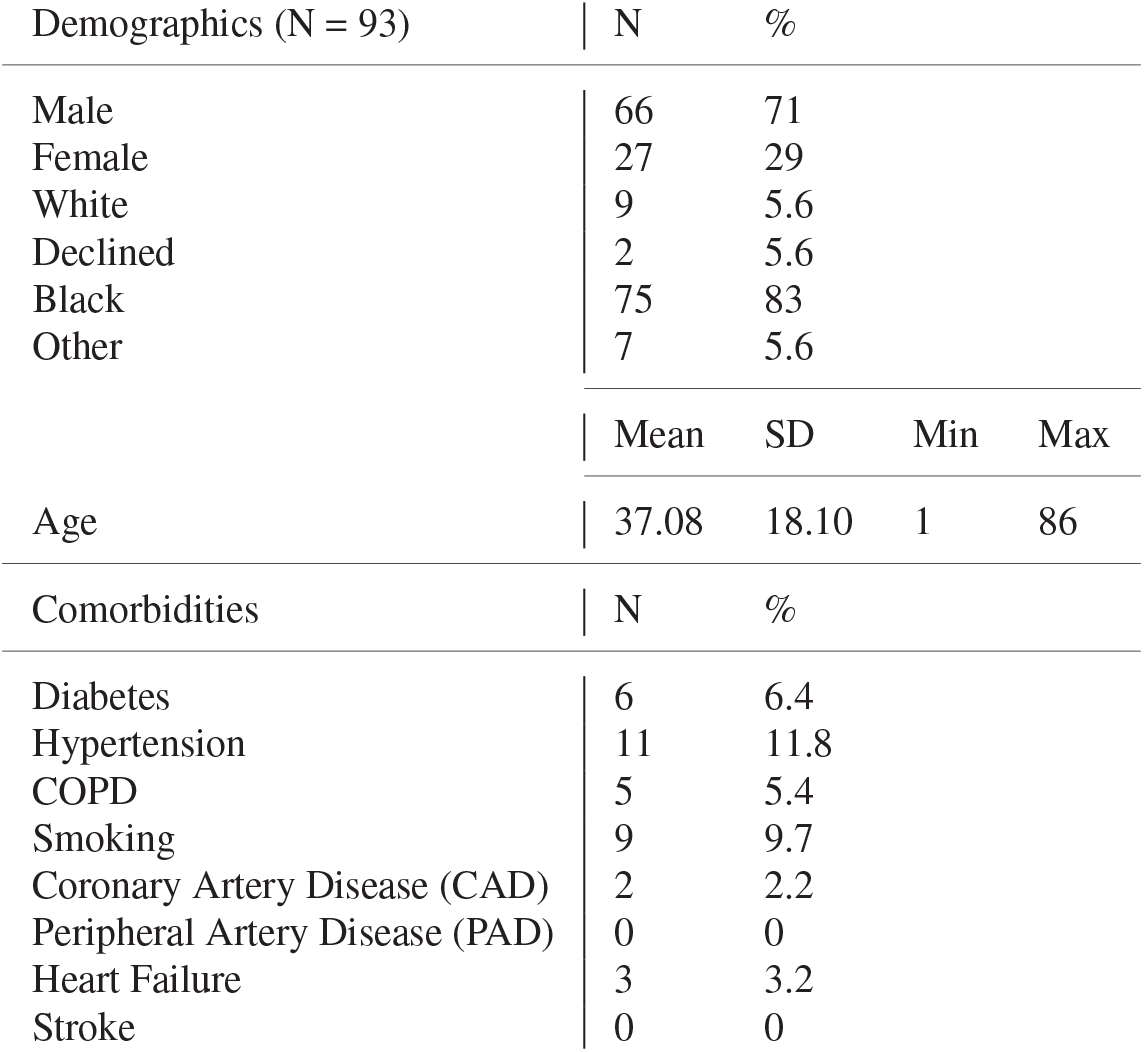
Non-Pathologic Aortic Cohort Patient Information

**Fig. 14.**
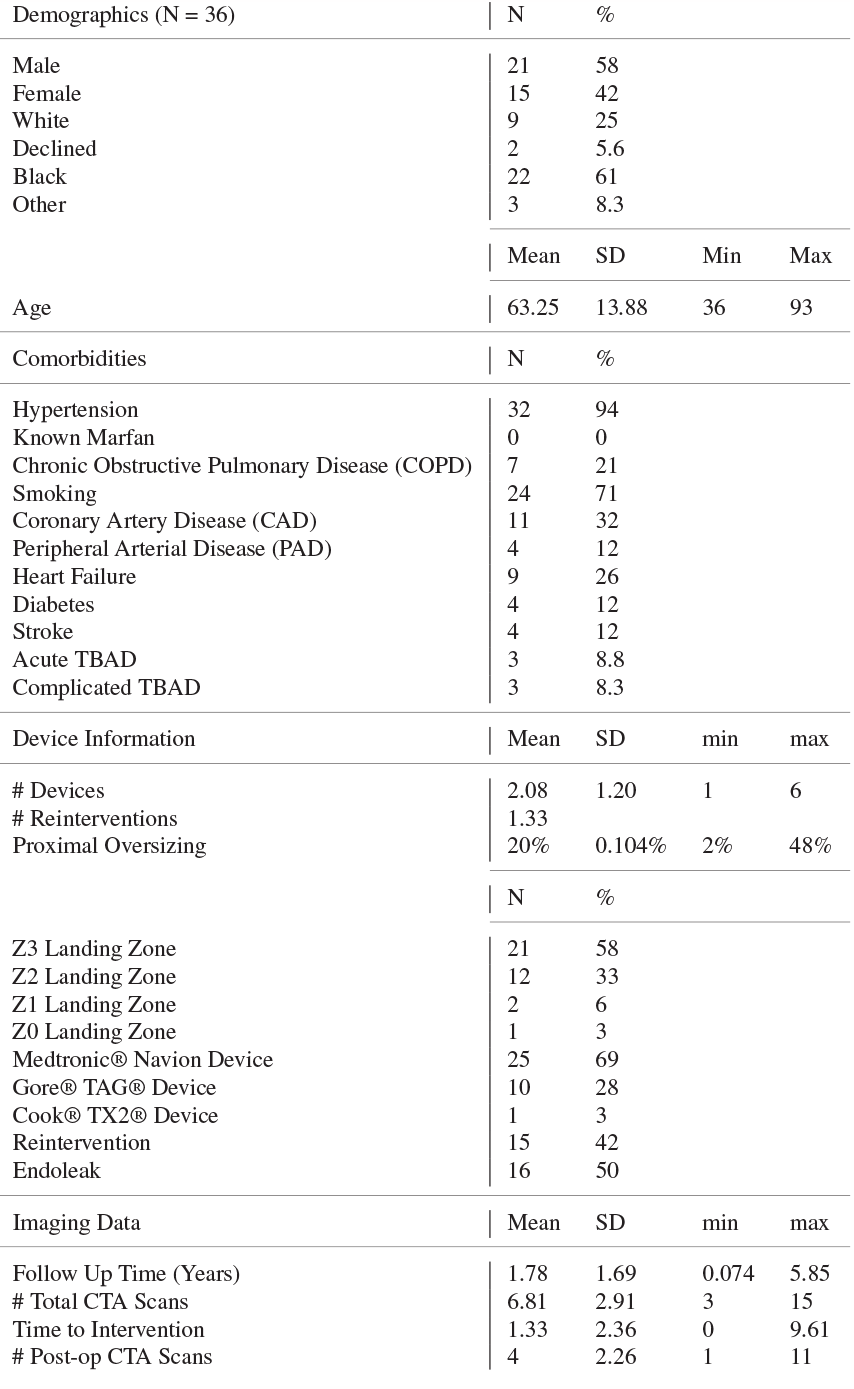
Type B Aortic Dissection Cohort Patient Information

Figure 11 shows how for the case of a small change in global size, as exhibited by the narrow range of the x-axis (9.5%), the fluctuation in Gaussian curvature (*δκ*_*g*_) accurately captures surface deformation just as well as the fluctuation in total curvature (*δ K*). This is to be expected as the main function of total curvature is to normalize size effects by mapping the aortic surface to **S**^2^. Thus, with little change in overall size, Gaussian curvature remains an effective indicator of shape. This is to be contrasted with what happens when size does change, as in the real patient data (400%) or in the idealized aorta setting shown in Figure 12 (50%). Gaussian curvature is no longer an effective indicator of shape when size also changes because it convolutes shape and size effects. On the other hand, total curvature normalizes size effects and effectively captures the degeneration in the shape of the ideal aorta. This experimental evidence in a controlled simulated system bolsters the empirical data and mathematical formulation, proving the significance of *δ K*.

## 4 Discussion

Type B aortic dissection (TBAD) is a life-threatening disease with significant associated morbidity and mortality [22–24]. While the old paradigm of open surgical repair was fraught with peri-operative risk, new minimally invasive approaches like TEVAR often trade a decrease in initial operative risk for higher risk of long-term repair failure. Proper identification of patients for TEVAR is therefore critical and necessitates the definition of an appropriate classification scheme [23, 26, 62, 63].

While previous work has focused on linking changes in aortic anatomy and suitability for repair, there remains a dire need to improve our understanding of how best to define geometric changes and to understand their impact on patient outcomes.

Projection of aortic anatomy into the (*K,ℓ*^−1^)-space provides an improved ability to differentiate aortas along the entire spectrum of growth and pathology, including both normal size-related development and pathologic shape-related changes. We demonstrate that in normal conditions, the aorta undergoes shape-invariant growth (see Figure 7), while, in diseased states, the aorta experiences shape fluctuations defined by increasing *δK*. As shown in Figure 10, *δK* significantly outperforms all other available measures of shape in predicting clinical treatment outcomes, including tortuosity, which is prevalent in the clinical arena.

Furthermore, because of the invariant global cylindrical geometry of the aorta, the parameterization of size is dependent only on the single length scale *ℓ*^-1^. Higher-dimensional characterizations of size, including area A_*T*_ and volume *V*, do not provide additional information [64, 65]. Thus, current efforts to replace 2*R*_*m*_ with area or volume [64, 65] are unlikely to yield substantially more information (Figures and 10). This universal size scaling provides the quantitative basis behind the utility of maximum diameter throughout the decades of aortic management and further validates the study of shape [63].

Figure 10 compares the efficacy of a single-variable space defined by maximum diameter (2*R*_*m*_), the clinical standard, with an enhanced shape-size feature space for predicting treatment outcomes. Using size as the sole metric of disease change, the cornerstone of imaging-based practice in other clinical contexts, is inherently problematic because a critical point between closely-spaced and overlapping populations is very sensitive to the available data. For instance, a well-known problem in *R*_*m*_-based criteria is the bias against smaller statured female patients because of the heavily male-weighted population-based statistics [66, 67]. This is best illustrated by Figure 10. While arbitrarily defined size classifiers did discriminate amongst normal aortas, successful TEVAR, and failed TEVAR, such scalars lack physical meaning and clinical generalizability outside of the specific cohort being analyzed. This is partly due to natural population-level variation in aortic size in addition to the inherently operator-dependent nature of aortic size measurements (such as diameter) [34].

The addition of the second axis (*δK*) alleviates this issue by providing a quantifiable and reproducible shape scalar. *δK* also captures the global geometry of the aorta and is operator independent. The combined 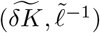-feature space demonstrates greater than 90% classification accuracy for the same three cohorts. The addition of a tangible shape axis also offers enhanced interpretability of the underlying geometric trends driving aortic pathology. Such a classification space can be clinically applied to pre-operative treatment planning for aortic dissection patients. *δK* outperforms previously described shape metrics in both the clinical and engineering literatures. Clinically-derived shape measures, such as the tortuosity index, are predominately acquired from aortic centerlines [25, 68]. These measures underperform compared to *δK* and are no better than *ℓ*^−1^ alone for characterizing aortic disease pathology from geometry. As such, future analysis of *δK* is liable to demonstrate substantial clinical application of *δK* as a clinical outcomes predictor.

No general theory exists on the meaning of *δK* divergence. In soft matter systems such as spherical vesicles, where size weakly changes, and in dynamic systems, rapid fluctuations of Gaussian curvature have been linked to so-called ‘topologic catastrophes’ indicative of physical instability [48, 49, 69]. While we do not inherently study aortic stability as it relates to clinical rupture, we show that aortas with high *δK* independently classify as high risk for clinical complications and poor outcomes. It is therefore reasonable to conjecture that vertical divergence in the 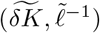-space is a sign of aortic instability and an indicator of suboptimal suitability for endovascular repair.

The morphologic evolution of biological structures is tightly integrated with disease development in many other contexts. *δK* is a size-independent shape metric, and because it only requires extrinsic geometric information, this procedure can be applied to any surface mesh geometry. For instance, 3D imaging is extensively used to analyze lung nodules for malignancy [1], breast lesions for tumor growth [70], liver irregularities for cirrhosis [2], cerebral aneurysms [71], and the left ventricle for heart failure [72].

As with the aorta, size-based criteria form the mainstay of clinical approaches, while shape is qualitatively used but has been proven difficult to quantify until now. This methodology is based on a general geometric and topologic foundation, and future analysis will be needed to validate its extension to other clinically-relevant problems of characterizing disease through analysis of shape change in medical imaging.

## A Appendices

### A.1 Demographic Information

### A.2 Aortic Segmentation and Post-Processing from CTA Imaging

#### A.2.1 Segmentation

Segmentation of the aorta is performed using a custom semi-automated workflow built to characterize global aortic geometry. The process first uses a threshold filling function to select voxels based on greyscale values. The threshold is then manually corrected by examining each axial slice, erasing non-aortic tissue, and performing other corrections as needed. The segmentation is cut at the aortic sinus, the branching of the brachiocephalic artery, and at the celiac artery. As the goal is to analyze aortic geometry, branch vessels are removed from the segmentation using a curved selection with 3D edit tool. Segmentation accuracy is independently reviewed by the senior author.

#### A.2.2 Noise Reduction

Segmentation of accurate patient-specific geometries from real CTA imaging data comes with several sources of noise, including inconsistent imaging quality and/or resolution, imaging artifacts, non-contrast-enhanced tissues like aortic thrombus, or residual variability between segmentations. Therefore, noise reduction is a key step of the workflow and includes the application of a thorough segmentation smoothing algorithm and a robust meshing algorithm. For each algorithm, key parameters are varied so that 15 distinct models are created for the one segmentation. This significantly reduces segmentation noise, as the models are aggregated later in the geometric analysis workflow.

#### A.2.3 Smoothing

The smoothing algorithm is a five-step process optimized to decrease noise while preserving local geometric features. It consists of dilation of the segmented geometry, the application of a mean filter, the application of a median filter, the application of a recursive Gaussian filter, and erosion. The segmentation is dilated (and later eroded) by four pixels. The mean and median filters compute each pixel’s value as a statistical mean and median of neighboring greyscale values within a radius of five pixels. The recursive Gaussian filter applies a smooth blur to the segmentation mask. Three models with three different widths of Gaussian kernels are made: a standard deviation of six, seven, and eight pixels in each direction for segmentations of dissections and four, five, and six pixels for segmentations of non-pathologic aortas. These parameters are the result of a thorough parameter sweep, in which the effect of changing each parameter on the resulting curvature values was analyzed.

#### A.2.4 Isolation of the Outer Surface

As aortas are systems of highly varying curvatures, a constant orientation of the normal vectors is maintained by isolating of the outer surface of the segmentation. The outer surface is isolated by making a closed surface of the outer shell of the aorta whose ends are defined by planes. The intersection of the planes defining the end of the segmentation and the shell creates rims of sharp surface curvature that are removed later in the workflow (see Appendix A.4).

#### A.2.5 Meshing

A triangular mesh for the outer surface is generated for each smoothed segmentation in ScanIP by creating a Matlab (2021b, Mathworks, Natick, MA) surface model with a minimum feature size of 0.5 mm and at 5 separate dimensionless coarseness parameters (on a scale from −50, very coarse, to 50, very fine, according to the program’s internal scaling): −35, −30, −25, −20, −15. This correlates to a mesh density of 0.5 elements/mm^2^ to 1.5 elements/mm^2^. These parameters represent optimal mesh densities and were found by studying the effect of changing the mesh density on calculated curvature values for analytically ideal shapes (e.g. a sphere, torus, cylinder). ScanIP uses a robust meshing algorithm, called “+FE Free,” to create accurate surface meshes for complex surface geometries. First, the algorithm generates a “+FE Grid” high quality mesh characterized by constant element edge lengths. Then, the surface is remeshed with the “+FE Free” algorithm with adaptive surface remeshing to enhance the mesh quality.

Therefore, for each segmentation, the 15 different outer surface meshes are created and analyzed as a result of the 3 smoothing parameter variants and 5 meshing parameter variants. This controls for the known variability of discrete derivatives calculated on meshed surfaces [50].

### A.3 Calculation of the Shape Operator

The Rusinkiewicz algorithm of estimating surface curvature computes the derivatives of surface normals [50] in order to calculate the per-vertex shape operator for each mesh. This algorithm excels at handling irregular triangular surface meshes, efficiently performing computations, and providing robust results on local surface neighborhoods. Briefly, the algorithm calculates the per-vertex shape operator as a weighted average of the shape operators of immediately adjacent faces. The per-face tensors are computed using a finite-difference approximation defined in terms of the directional derivative of the surface normal [50]. The algorithm was implemented in MATLAB [73].

First, the per-face normal vectors are calculated as 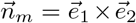, in which 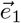 and 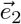 are two unit-length edges of the triangular surface mesh. The per-vertex normal vectors 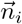 are then calculated using a weighting algorithm to accurately estimate vertex normals [74]. For each face *m* of area *A*_*m*_ containing three edges 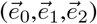, each of the 3 vertices is assigned a weight: 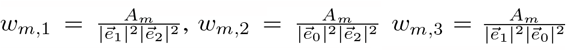. Then, the contribution of face *m* to the normal vector of vertex *i* is 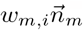 h is added to 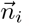. After this is done for all faces containing vertex *i*, the per-vertex normal vectors are normalized to unit length [50].

A per-vertex 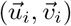 coordinate system is calculated in which 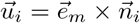 and 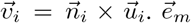 indicates a face edge vector touching vertex *i*. This per-vertex coordinate system will later be utilized for averaging the per-face shape operator with the contributions of adjacent faces to calculate the per-vertex tensor [50].

Next, the per-face shape operator is calculated. For each face *m*, a 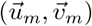 coordinate system is defined as 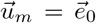 and 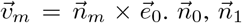, and 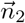 are defined as the normal vectors of the three vertices constituting the face. Then, the second fundamental form II is calculated by solving a least-squares problem:

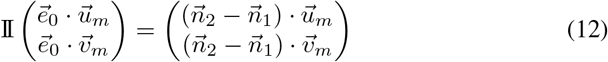

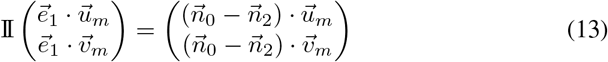

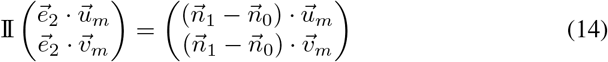

The shape operator is then projected onto each of the face’s vertices. To avoid a “loss” of curvature at the change from the vertex-based coordinates 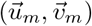 to the face-based coordinates 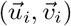), one of the coordinate systems is rotated to be coplanar with the other, as previously performed in the literature [50]. Similarly to the calculation of the per-vertex normal vectors, the contribution of each touching face is weighted to form the per-vertex shape operator. In contrast to the weighing procedure for calculating the normal vectors, previous literature [50**?**] utilizes a “Voronoi area” weighting procedure for finding the per-vertex shape operator, which is followed here. This calculates 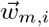 as the portion of the area of face m that lies closest to vertex *i*. This approach has been found to produce the best estimates of curvature for faces of varying sizes and shapes [50]. This tensor is then diagonalized to obtain the per-vertex principal curvatures *k*_1*i*_, *k*_2*i*_. This results in the per-vertex principal curvatures for each aortic surface geometry’s 15 meshes.

### A.4 Artifact Removal

The isolation of the outer surface of the aortic segmentations, which is performed to maintain a consistent orientation of normal vectors, results in the creation of flat edges with rims of sharp curvature. The contribution of these flat edge regions are removed before calculating the curvature functions. As artifacts, these points have outlying curvature values. Flat edges are removed by removing vertices with 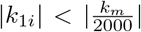 and 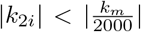, where *k*_*m*_ is the average of the per-vertex mean curvature (*k*_*m*_ = (*k*_1*i*_ + *k*_2*i*_)/2) for the entire surface. This threshold value was carefully chosen to be overly specific, and each segmentation is manually verified to confirm that only the flat edges are removed.

Removing the sharp rims required separate criteria for normal and diseased aortas. For the diseased aortas, points with 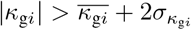 and 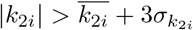 are removed. Here, 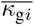 indicates the standard deviation of (*κ*_*gi*_) of the aorta, and 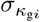 indicates the standard deviation of the Gaussian curvature. Similarly, 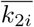is the mean second principal curvature (*k*_2*i*_), which is defined as the principal curvature with a larger absolute value. For normal aortas, points with 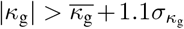 and 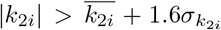 are removed. These parameters are manually chosen to remove outlying points without removing points actually on the outer surface. On average, around 1 −2% of mesh vertices are removed as part of the rims of sharp curvature. Representative examples of the results of both artifact removal procedures are shown in figure 15. For the ideal shapes, no points are removed for the sphere, points of positive Gaussian curvature are removed for the catenoid and pseudosphere, and the same procedure as for diseased aortas is utilized for the torus and cylinder, resulting in an average of 0.03% of points being removed.

**Fig. 15.**
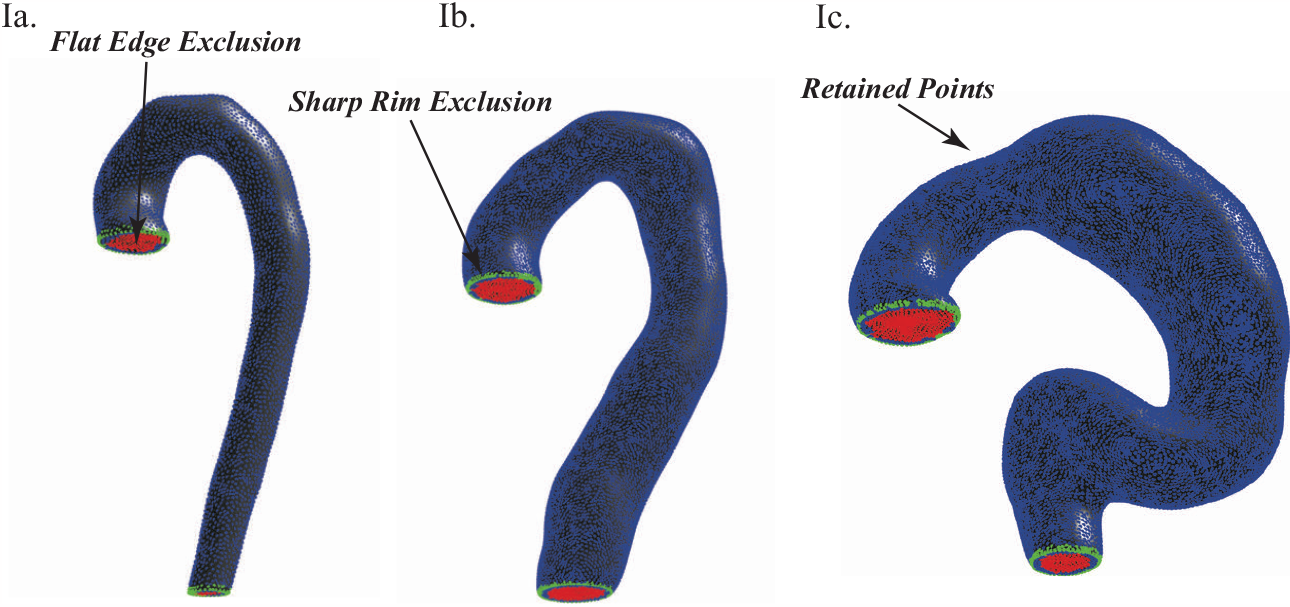
Demonstration of artifact removal procedures. Points excluded using the flat edge removal procedure are shown in red, and points excluded using the sharp rim removal procedure are shown in green. Retained points are shown in blue. Representative examples are shown for Ia. a non-pathologic aorta, Ib. a diseased aorta in the successful TEVAR cohort, and Ic. a diseased aorta in the failed TEVAR cohort. As can be seen, these artifact removal procedures are more specific than sensitive, in that not all artifact points have been removed, but very few (if any) non-artifact points are incorrectly excluded. Images are not drawn to scale.

### A.5 Jensen-Shannon Divergence of Within-Partition Gaussian Curvature

The method requires that *κ*_*gi*_ is relatively constant inside each surface partition, such that 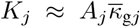, where 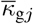 is the mean Gaussian curvature within partition *j*. As discussed in section 2.3.2, the partitioning is implemented via a simple Voronoi decomposition without explicitly taking local curvatures into account. To calculate the degree of *κ*_*gi*_ variability within each partition for a given decomposition, the Jensen-Shannon Divergence (JSD) is used. The JSD is an information-theoretic measure of dissimilarity among probability distributions, and it is used here to quantify the discrepancy between the ideal partition of weakly varying *κ*_*gi*_ and the true surface partition [76].

First, an aortic surface with weakly varying *κ*_*gi*_ is defined as having a *κ*_*gi*_ distribution centered at 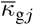, the mean Gaussian curvature in the partition, with a standard deviation of 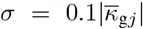. Although this definition is arbitrary, it precisely defines a tight distribution with minimal variation in *κ*_*gi*_ that appropriately scales with the magnitude of curvature in a partition. For distributions *f*_1_(X) and 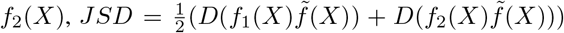, where 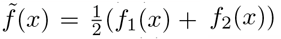is the common distribution and 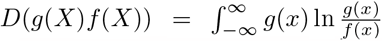 is the relative entropy, which captures the divergence from the given probability distribution *f*(*X*) *t* the reference distribution *g*(*X*) [77, 78]. The discrete form 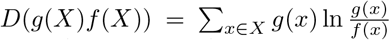 is calculated. The Freedman–Diaconis rule is used to optimize the calculation of the *JSD* by properly discretizing the *κ*_*g*_ distribution *f*(*X*) for partition-level data 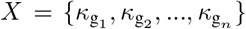 into m bins, where 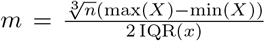 [79]. The reference distribution *g*(*X*) is defined by randomly sampling 10,000 values from a Gaussian distribution with a mean of 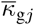 and 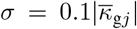 and then creating a histogram. Both distributions are normalized to probability mass functions prior to calculating the JSD of the partition.

Figure 16 visually demonstrates the calculation of the Jensen-Shannon divergence. On average, partitions have a JSD of under 0.2 (see dotted line in panel III), compared to a maximum bound of ln(2). This demonstrates that partitions have a relatively weakly varying *κ*_*g*_ when *k* = *A*_*T*_ /*ℓ*^*−*2^ partitions are defined. Figure 16III also shows that the JSD increases for a coarser partitioning (*k*/10 partitions). In this limit, the method of calculating *K* is calculation of is reduced for a higher density partitioning (*k* ∗10 shown). This identifies a range of partition densities within which the extrinsic calculation of local *K* is valid.

**Fig. 16.**
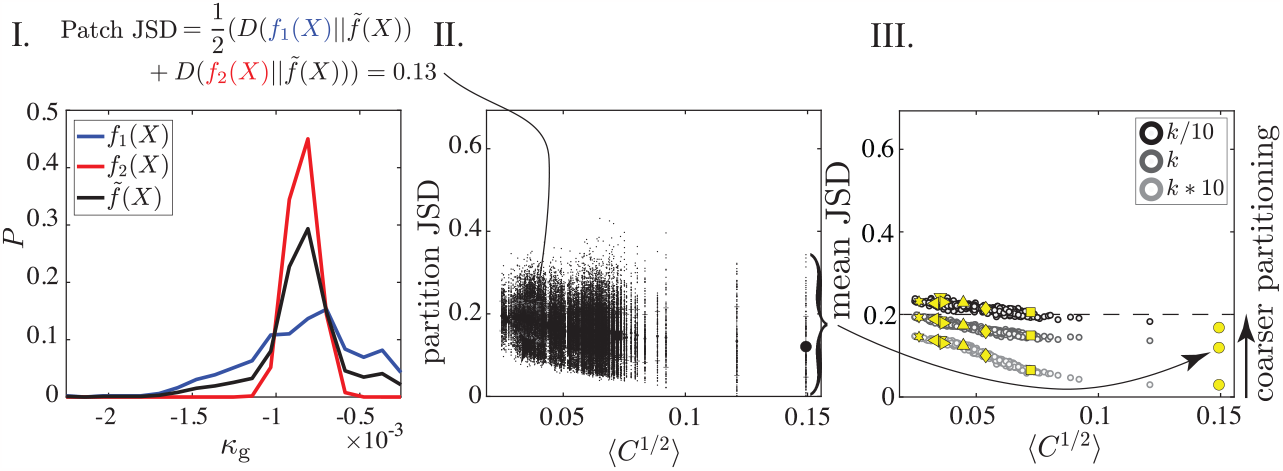
Calculation of Jensen–Shannon divergence (JSD) to quantify the variability of within-partition Gaussian curvature. I. The JSD is calculated for a partition by comparing *f*_1_(*X*), the *κ*_g_ distribution, with *f*_2_(*X*), the reference distribution with weakly-varying *κ*_g_. II. Plot of the JSD for all partitions versus ⟨*C*^1*/*2^⟩ for the corresponding aorta. III. The mean JSD is calculated for all 302 aortas (yellow symbols correspond to the canonical 8 aortas, see figure 1). The JSD is compared for three partitioning densities. At the base level of *k*, JSD *<* 0.2 out of a theoretical maximum of ln(2). When *k* increases by a factor of 10, the mean JSD decreases for all patients, indicating that the weakly-varying curvature partition assumption becomes more valid. When *k* decreases by a factor of 10, the mean JSD increases for all patients, indicating that the weakly varying *κ*_g_ assumption is less valid.

### A.6 Sensitivity to Partition Size

As detailed in section 2.3.2, each aorta is subdivided into *k* = *A*_*T*_ *ℓ*^*−*2^ partitions using a *k*-means algorithm. This achieves an approximately constant number of partitions, each sized according to a self-contained length scale, across all aortas. Figure 16III shows that JSD < 0.2 and the extrinsic computation of *k* remains valid for number of partitions greater than *k*. If the number of partitions decreases, JSD increases and the computation of *K* is invalid. In this section, the impact of partition number on the curvature functions *δK* and ⟨*C*^1*/*2^⟩ is studied. The number of partitions is increased or decreased by a factor of 10. The calculations are also performed on the level of a single mesh element.

Figure 17 shows the data presented in Figure 7III of the main paper with the calculations performed on different partition densities and with no partitioning at the level of the mesh. As outlined in the main text, *δK* captures the balance of positive and negative curvatures across the aortic surface in a globally scale-invariant manner. For aortas of different size but constant shape (due to normal growth during development), *δK* must be constant. Figure 17IV shows that *δK* does not follow the trend when the computations are performed on the level of the mesh. We hypothesize that the breakdown of the computation in this limit occurs because the mesh is composed of flat Euclidean triangles. Diguet’s Theorem states that for surface patches of con-stant *k*_*g*_ (low *JSD* in our case), the surface area scales as 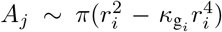 where *r*_*i*_ extends from the patch center along a geodesic line [80, 81]. As said, for 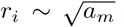, the area calculation does not correctly account for the local curvature and one would need access to the metric to perform the integration. The breakdown of the scaling with coarsened partitions, shown in Figure 17-I, is also discussed in Appendix A.5; *JSD* > 0.2 invalidates the ability to take *k*_*g*_ outside the integral.

**Fig. 17.**
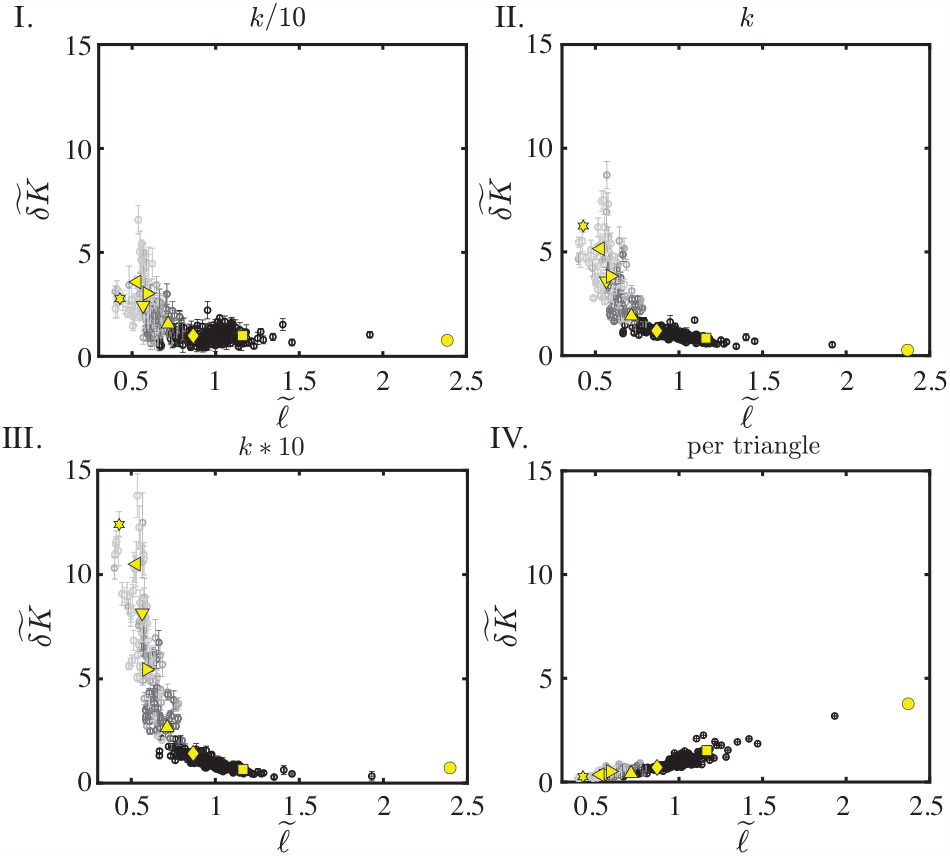
Tuning the partition number *k* demonstrates the sensitivity of the method to partition size. The surface is partitioned into *k* = *A*_*T*_*ℓ* ^*−*2^ partitions to obtain *A*_*j*_, the area over which the partition-level total curvature 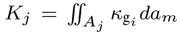 is taken. This number remains on the order of *O*(100) for many aortic shapes and sizes. I. When *k* is divided by 10, there is a clear increase in noise causing higher variation in *bK* for the normal aortas, poor separation of the three patient groups, and a suppressed divergence in *bK* for highly diseased aortas (light gray). II. The base level of *k*. III. Multiplying *k* by 10 (resulting in a larger partition number and smaller partition size) shows that the general feature signal contained in *bK* is retained. IV. Performing the calculation at a per-triangle level, at which 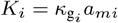 is the total curvature of one triangle, inverts the direction of the trend and degrades the signal. This is because individual element areas *a*_*m*_ are locally Euclidean and thus cannot individually approximate a curved surface

### A.7 Ideal Shapes

Before continuing with the geometric analysis workflow, the ideal shapes used in Figure 7II and 7III in the main text are created to demonstrate that the sum of total curvature (Σ*K*) is a topologic invariant. Spheres, tori, cylinders, pseudospheres, and catenoids of varying sizes are defined using mATLAB. For each geometry, point clouds are defined from the respective functional form and triangulated into a surface mesh. Cylinders are created with radii *r* = {5, 7, 9, 14, 28, 35} mm and heights *h* = 2*r* for a total of 6 models. Spheres are defined with radii *r* = 7, 8, 9, 10, 13, 20, 25, 50 mm to obtain 8 models. Tori are defined using the following parametric equations:

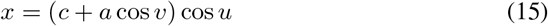

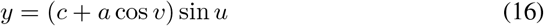

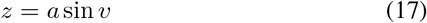

for *u, v* ∈ [0, 2 *π*), where *c* = {11, 14, 20, 35, 50, 60} mm and *a* = *c*/2 (for 6 models).

Likewise, the following Cartesian parametric equations define the catenoids:

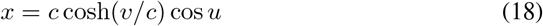

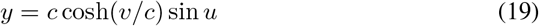

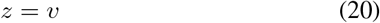

for *u* ∈ [0, 2 *π*), with *c* = {2, 8, 10} mm and *v* ∈ [−*c, c*].

The pseudosphere is similarly defined according to the Cartesian parametric equations

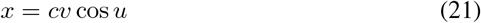

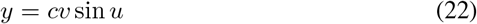

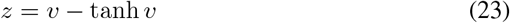

for *u* ∈ [0, 2 *π*), with *v* in *v* = 5 tanh (0.5*x*) mm (*x* ∈ [−*π, π*)) and *c* = {25, 40, 50, 75, 100} mm.

These specific parameters for the ideal geometries are chosen to obtain shapes at a wide range of sizes. The same algorithm as for the aortas is used to compute the per-vertex shape operators.

### A.8 Other Shape Metrics

Figure 18 analyzes previous shape metrics from the literature. In addition to quantifying rupture risk using Gaussian curvature, investigators predicting aneurysmal rupture risk have utilized other geometric indices, including the L2-norm of the Gaussian curvature 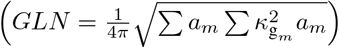, area-averaged Gaussian c ture 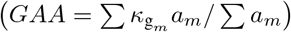, area-averaged mean curvature 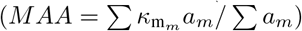, and L2-norm of the mean curvature 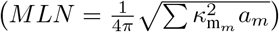 [82–84]. While one study concluded that the GAA is highly correlated with wall stress [85], another concluded that the GLN and MLN best classified unruptured vs. ruptured AAAs [86].

**Fig. 18.**
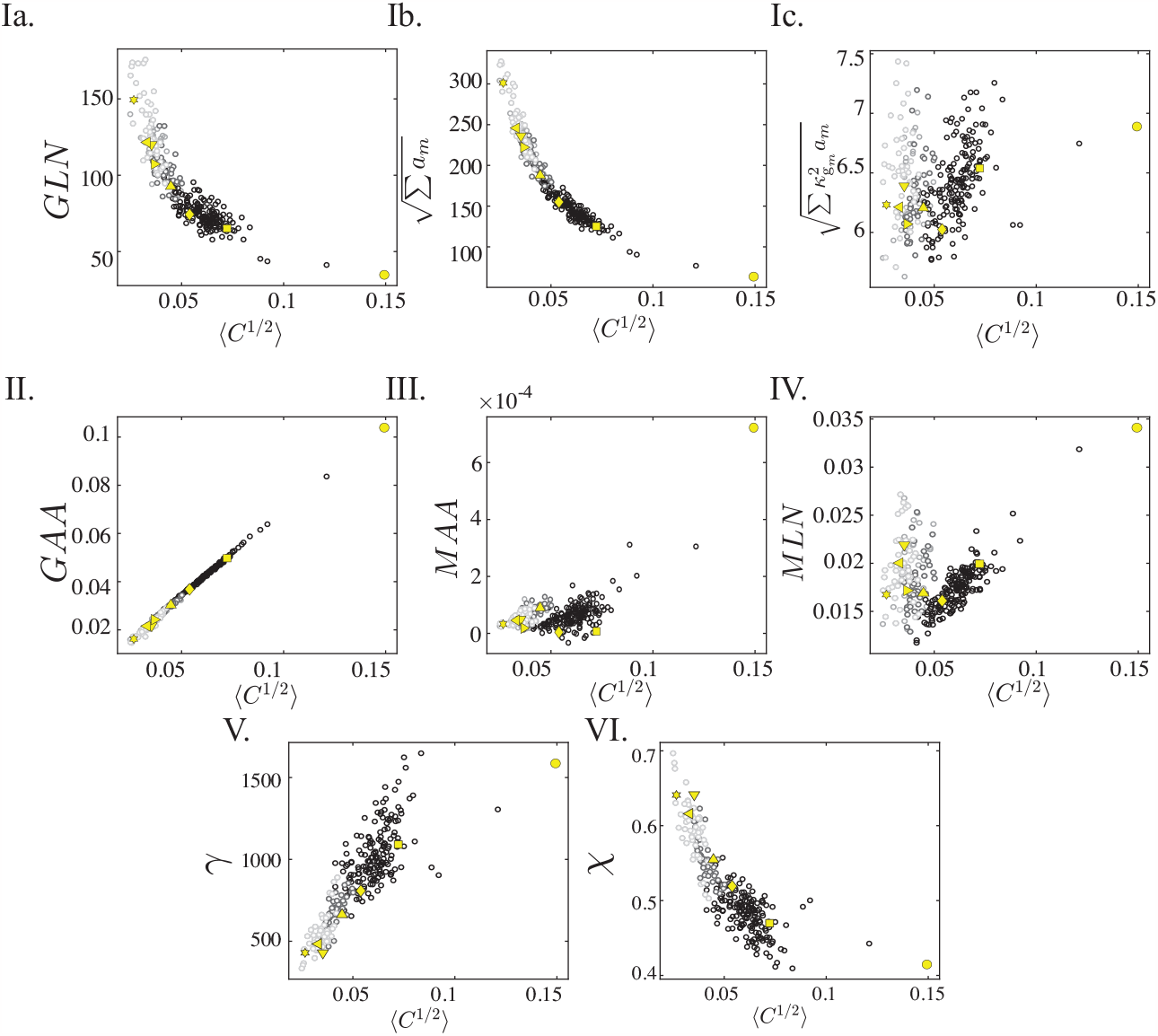
Shape metrics from the literature. Ia. The GLN distinguishes the patient groups, but it scales with ⟨*C*^1*/*2^ ⟩, as can be seen from its increasing value with increasing size of the non-pathologic aortas (while *bK* remains constant throughout this regime). The GLN is mathematically composed of two parts. Ib. The first term is simply the sum of the individual element areas, which is the total aortic area. Ic. The second term 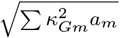 carries no information, as evidenced by the absence of a trend. II. The GAA is a size metric as it linearly scales with ⟨*C*^1*/*2^ ⟩. III. The MAA carries no meaningful information, as it cannot separate normal from seased aortas IV. The MLN similarly does not correlate with aortic disease state. V. The flatness index (*γ* = *A*^3^*/V* ^2^) s les with size an is non-constant for non-pathologic aortas of different sizes. VI. The sphericity index 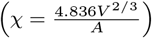 also scales with size.

The drawbacks of these measures become evident when applied to the patient set in figure 18. Panel I demonstrates how the GLN is essentially a reparameterized version of *A*_*T*_, as it clearly scales with size, with the one term being mathematically equivalent to *A*_*T*_ (Ib) and the second term demonstrating no correlation with the data (Ic). The GAA (panel II) again scales with size, and the MAA and MLN are poor classifiers of aortic disease state (panels III and IV). Panels V and VI test shape measures from the broader biological literature that examine the scaling of areas and volumes: the flatness index *γ* = *A*^3^/*V* ^2^ and the sphericity index 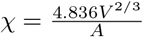 [87]. The sphericity index is defined as unity for a sphere such that 0 < χ < 1, making it a bounded form of the flatness index. The two measures are related by *γ* ∝ χ^*-*3^. As shown, both measures simply scale with size. Neither provides a constant baseline for the non-pathologic patients.

### A.9 Finite Element Simulations

#### A.9.1 Simulation of a Sphere

A sphere is defined with inner radius 3 mm and thickness 0.1 mm (see figure 11). A linear tetrahedral mesh is applied with a target of 5 elements through the thickness for a total of 214,643 elements. A neo-Hookean material model is used with material parameters 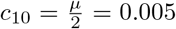 MPa and *D* = 2 MPa (*μ* is the shear modulus and *D* is the bulk modulus). The inner surface is pressurized to 30 mmHg. During the pressurization process, no growth is prescribed to the aortic wall. This is followed by growth in randomly selected surface partitions at a rate of 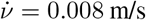. Pressurization is performed for 10 ms and growth for 50 ms for 60 ms total in the simulation. No boundary conditions are applied. Eighty frames are selected from the simulation. To obtain the consistent orientation of normal vectors, the outer surface is isolated. The surface is smoothed to reduce simulation noise, and the geometry is re-meshed in ScanIP to apply an optimal mesh for calculating discrete derivatives.

#### A.9.2 Simulation of an Idealized Aorta

To demonstrate the loss in utility for Gaussian curvature when global size significantly changes, pressurization and growth are modeled in an idealized aortic geometry (see figure 12). An ideal aorta is defined in the shape of a candy cane consisting of a cylinder of length 140 mm attached to a half-torus with a cross-sectional radius of 14 mm and a toroidal radius of 28 mm. A linear tetrahedral mesh with a target of 5 elements through the thickness for a total of 239,777 elements is used. The ideal aorta is simulated using the same material property as the sphere but with slightly different pressurization and growth parameters. The simulation applies 225 mmHg of pressure in 40 ms followed by 60 ms of growth in randomly selected surface partitions at a rate of 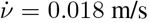., resulting in 100 ms of simulation time in total. The surface partitions are manually selected to be roughly evenly distributed along the surface. From the simulation, 100 frames are selected. The outer surface is extracted to obtain a consistent orientation of normal vectors. Then, the geometry is smoothed to reduce simulation noise, and it is re-meshed in ScanIP to apply an optimal mesh for calculating discrete derivatives.

## Acknowledgments

We acknowledge the support of the National Institutes of Health, NHLBI, R01HL159205. The Center for Research Informatics is funded by the Biological Science Division at The University of Chicago with additional funding provided by the Institute for Translational Medicine, CTSA grant number ULITR000430 from NIH. We are grateful to Enrique Cerda, Efi Efrati, Haim Diamant, and Thomas Witten for careful reading of the text and detailed comments.

